# Dual roles of DNA methylation in gene expression and chromatin structure in a stick insect

**DOI:** 10.1101/2025.10.30.682900

**Authors:** Nicholas P. Planidin, Clarissa F. de Carvalho, Zachariah Gompert, Patrik Nosil

**Affiliations:** Centre d’Écologie Fonctionnelle et Évolutive (CEFE), Université de Montpellier, CNRS, EPHE, IRD; Montpellier, 34293, France; Station d’Écologie Théorique et Expérimentale (SETE), CNRS; Moulis, 09200, France; Departamento de Ecologia e Biologia Evolutiva, UNIFESP; Diadema, 09972-270, Brazil; Department of Biology, Utah State University; Logan, Utah, 84322, USA

## Abstract

Gene body methylation is one of the most taxonomically widespread epigenetic marks, yet its function and evolutionary history are poorly understood. While there is a positive association between gene body methylation and gene expression in many organisms, the mechanisms behind this relationship are unclear. Here, we investigate the function of gene body methylation in the stick insect *Timema cristinae*. We compare patterns of genome-wide gene body methylation to gene expression, open chromatin peaks, and chromatin compartments. We find that 95% of genes with gene body methylation occur in open chromatin compartments (euchromatin). Furthermore, highly expressed genes are impoverished in methylation at their transcription start site (TSS), which is associated with a peak of open chromatin. These findings suggest that the positive correlation between gene body methylation and gene expression is due to chromatin compartment structure. Yet, methylation around the TSS is associated with gene expression, possibly playing an inhibitory role as in vertebrates. Lastly, comparative analysis of *Apis mellifera* reveals a similar relationship between methylation and chromatin compartments, but not gene expression. These results suggest a possible explanation for the heterogeneous association between gene body methylation and gene expression and give insight into DNA methylations role in regulating gene expression.

## Introduction

Genomes produce phenotypes through the expression of genes. Natural selection, in turn, acts on phenotypes, allowing evolution to shape genetic variation. It is therefore essential to understand the mechanisms regulating gene expression, to fully understand evolutionary dynamics. One of the primary mechanisms regulating gene expression is epigenetic marks, molecular markers associated with DNA that remain stable across cell divisions ^1^. Through changes in the pattern of epigenetic marks, a single genome can produce a diversity of cell types, develop complex phenotypes, and respond to environmental stimuli ^2^. Thus, understanding the function and evolutionary history of different types of epigenetic marks can give key insight into how gene expression is regulated and how such regulation evolved.

DNA methylation (5-methylcytosine) is one of the most functionally diverse and well-studied epigenetic marks (Fig. 1A). In plants and fungi, DNA methylation occurs at a variety of genomic motifs (CpG, CHG and CHH; H = A, T or C) and occurs most frequently in repetitive elements and transposons^3,4^. In contrast, in animals DNA methylation is largely constrained to symmetric CpG dinucleotides^5^. Furthermore, in vertebrates most CpG sites are highly methylated with the exception of active promoters^5^, whereas in invertebrates most CpG sites lack methylation except for the body of certain genes^6,7^. There are also a number of species which completely lack DNA methylation, including the model invertebrates *Drosophila melanogaster* and *Caenorhabditis elegans*^8^. Such variation in DNA methylation makes it clear that it has undergone a variety of gains and losses of functions across Eukaryota, leading to debate over which functions are ancestral versus derived^9–11^.

**Figure 1.**
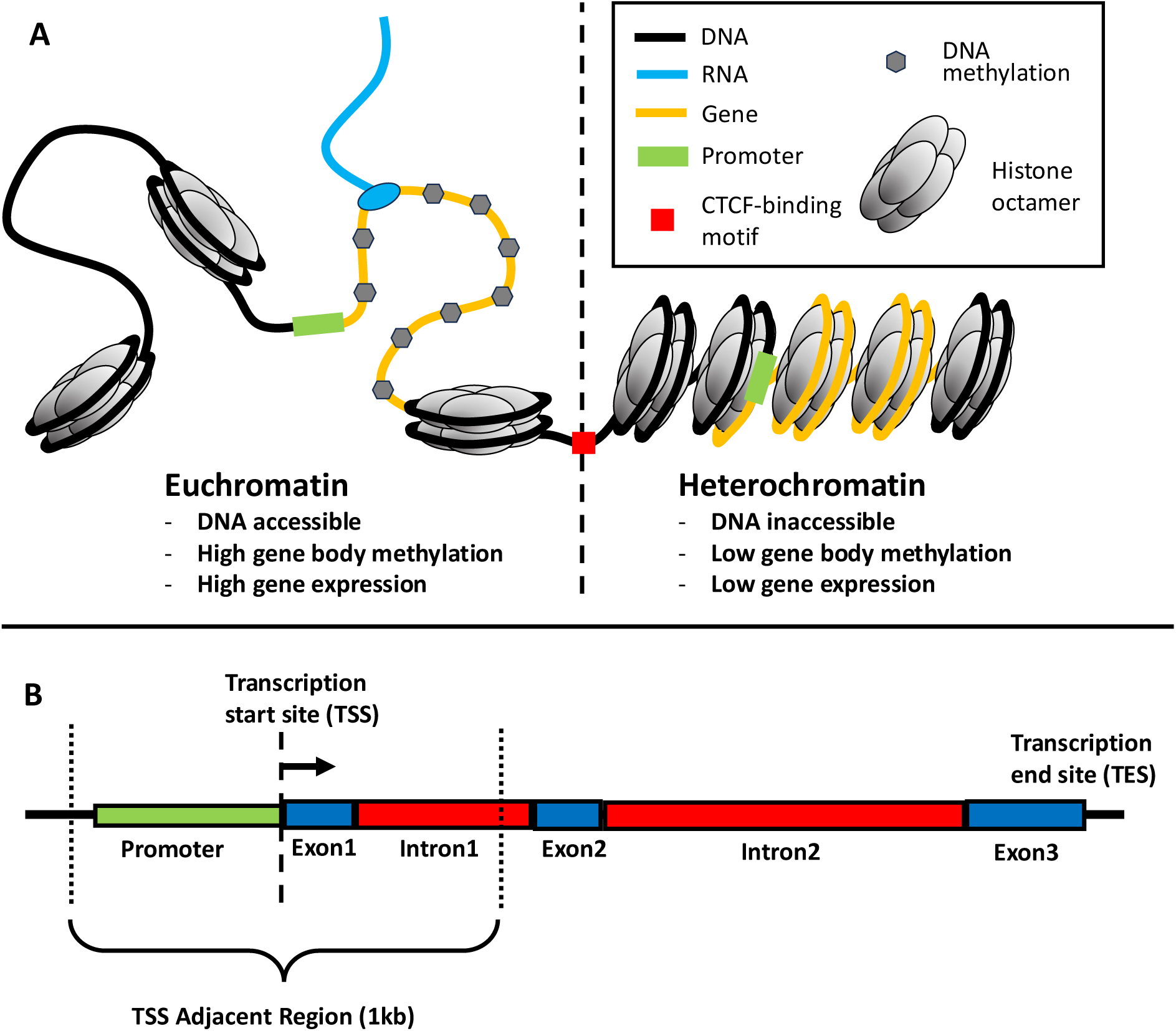
An overview of epigenetic marks examined in this study. (A) A schematic of the patterns of epigenetic marks in invertebrates, for the epigenetic marks we analyze in this study. CTCF stands for CCCTC-binding factor. (B) A schematic of gene structure and how it relates to the gene body regions analysed in this study.

Despite the fact that gene body methylation is taxonomically widespread, its function remains poorly understood. An association between gene body methylation and greater gene expression has been observed in vertebrates^12^, invertebrates^11,13–15^, and plants^16–18^. However, there is evidence of gene body methylation having no association or a negative association with gene expression in all three groups^19–21^. This varying evidence has led to the development of many hypotheses for the function of gene body methylation, such as the regulation of housekeeping genes^7,16^, the stabilization of transcriptional noise^22,23^, or the control of alternative splicing^24–26^. DNA methylation is also mutagenic, as 5-methylcytosine mutates into thymine at an elevated rate^27^. It is therefore peculiar that highly methylated genes typically have a lower rate of genetic divergence^6,28,29^, suggesting that the stabilizing function of gene body methylation outweighs its mutagenic effects. There is ample evidence that functional gene body methylation is taxonomically widespread. However, its exact function and its consistency across the tree of life is debated^30^.

One way that DNA methylation regulates gene expression is through its effects on chromatin: the complex of DNA, RNA and proteins that determines the physical structure of eukaryotic genomes^31^ (Fig. 1A). At the smallest scale, DNA is packaged into nucleosomes, segments of ∼150 bp of DNA wrapped around an octamer of histone proteins (Fig. 1A). Through either packing or unpacking DNA into nucleosomes, segments of DNA can either be made accessible or inaccessible to other proteins, modifying its function. For example, if the transcription start site (TSS) of a gene is free of nucleosomes, it is more accessible to transcription factors, promoting gene expression^32^ (Fig. 1B). Thus, by stabilizing nucleosomes at the TSS, DNA methylation at promoters can suppress gene expression^33,34^. At the largest scale, chromatin can be categorized into two functionally distinct types; tightly packed and inactive heterochromatin, and loosely spaced and active euchromatin (Fig. 1A). As these two chromatin states physically separate within the nucleus, they form compartments the size of hundreds of kilobases to megabases, often called A (euchromatin) and B (heterochromatin) compartments^35^. DNA methylation also influences chromatin compartment formation by interacting with multiple histone modifications to condense and stabilize regions of heterochromatin^36^. Lastly, within a single chromatin compartment, distant genomic elements such as enhancers, promoters and co-expressed genes are brought into physical proximity and co-regulated within topologically associated domains (TADs)^37^. The formation of TADs is regulated by CCCTC-binding factors (CTCF), whose activity may be impacted by DNA methylation at CTCF-binding motifs^38,39^ (Fig. 1A). While DNA methylation is involved in the determination of chromatin structure at multiple scales, its relative importance in such processes and their effects on gene expression are still poorly understood outside of laboratory models such as human cell cultures, *Mus musculus*, and *Arabidopsis thaliana*.

As with other functions of DNA methylation, its interaction with chromatin is also diverse among taxa. While increased DNA methylation is associated with heterochromatin in most eukaryotes^40,41^, the opposite trend has been observed in electrophoretic studies of insects^42,43^. Furthermore, while the regulation of gene expression by promoter methylation is widespread among vertebrates and plants, this function seems to be lacking in many invertebrates and fungi^3,44,45^. Therefore, by studying the relationship between DNA methylation, chromatin, and gene expression together in a stick insect, we can better understand transitions in the function of DNA methylation during evolutionary history.

Here we use *Timema cristinae* stick insects^46^, an emerging model species for molecular evolution with relatively high levels of gene body methylation, to examine the function of DNA methylation. *Timema cristinae* is a flightless phytophagous stick insect found in the chaparral habitat of the Santa Ynez mountains of California. There is a long history of using *T. cristinae* to study adaptation and speciation^47–50^. As part of this work, a diverse molecular toolkit has been developed to study *T. cristinae*. Previous studies of *T. cristinae* have described its DNA methylation; showing that it differs between ecotypes, correlates with gene expression, and plays a role in genetic divergence among populations^51,52^. Furthermore, recently published haplotype phased genome assemblies for *T. cristinae* have used high-throughput chromatin conformation capture (Hi-C)^53^, providing the data necessary to investigate chromatin compartmentalization in this species.

In this study, we use the available molecular toolkit of *T. cristinae* in addition to newly generated chromatin accessibility (ATAC-seq) data and additional Methyl-seq data to investigate the function of gene body methylation in this species. We do so by comparing profiles of DNA methylation, nucleosome-free regions (open chromatin peaks), gene expression, and chromatin compartments across the genome. Therein, we test two hypotheses about the function of DNA methylation. First, whether methylation at the TSS is positively or negatively associated with gene expression (Fig. 2A). Second, whether DNA methylation is associated with open or closed chromatin (Fig. 2B). To conclude, we compare our findings in *T. cristinae* to the European honey bee, *Apis mellifera*, using publicly available data. Notably, our study is correlational and does not test the causal relationship underlying these associations. However, by examining the pattern of DNA methylation, chromatin state, and gene expression in insects, we gain insight into the evolutionary history of DNA methylation and its association with gene expression.

**Figure 2.**
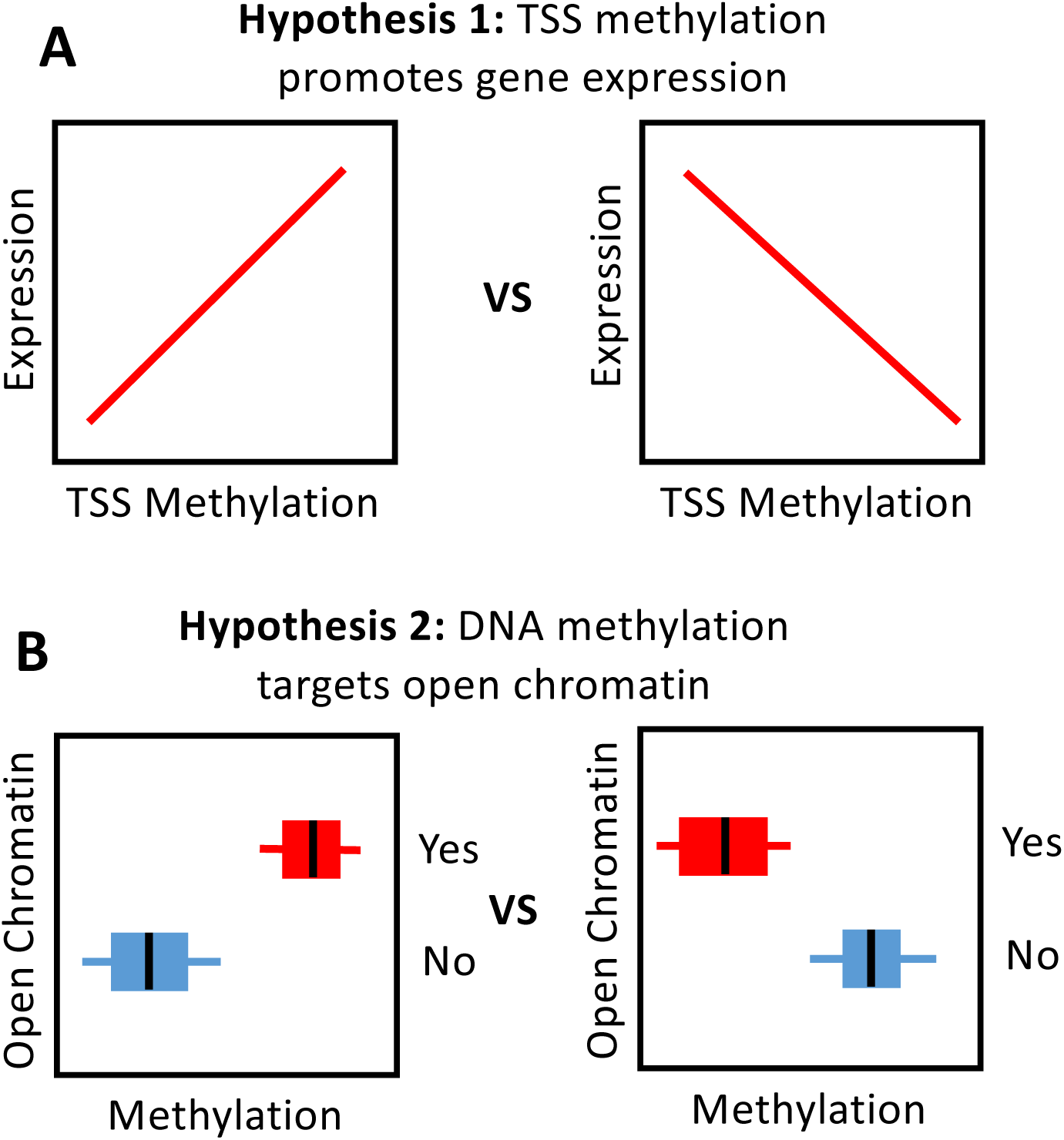
Hypotheses tested in this study. (A) The expected relationship between transcription start site (TSS) methylation and gene expression, depending on the function of DNA methylation. (B) Boxplots showing the expected relationship between DNA methylation and regions of open chromatin, depending on the function of DNA methylation. Boxplots display median (center line), quartiles (box), and quartiles plus 1.5 times the interquartile range (whiskers).

## Results

### TSS versus gene body methylation

It has been observed that promoter methylation negatively correlates with gene expression in some invertebrates^6,14,54^. Furthermore, in some model species, gene body methylation at the first exon and intron also have a negative relationship with gene expression^11,19,55^. So, we began our investigation by examining how the position of DNA methylation along the gene body relates to gene expression in *T. cristinae*. To do so, we measured DNA methylation and open chromatin peaks in 100 bp bins along the putative promoter region (here defined as the 500 bp upstream of the TSS as by Keller et al.^6^) and gene body, and compared them between highly expressed and unexpressed genes (here defined as the upper 95^th^ versus the lower 5^th^ quantiles of gene expression).

Methylation at the TSS and the gene body had opposite correlations with gene expression. In highly expressed genes, the region adjacent to the TSS (the promoter and 500 bp downstream of the TSS) had less methylation than unexpressed genes (4.5% versus 10% methylation, respectively; Wilcoxon rank-sum test: n_1_ = 315, n_2_ = 1107, W = 219476, p = 2×10^-12^; Fig. 3A; Fig. S1A). The TSS was also more likely to contain a peak of open chromatin in highly expressed genes (88% of TSSs with an open chromatin peak versus 23% in unexpressed genes; Wilcoxon rank-sum test: n_1_ = 315, n_2_ = 1139, W = 62809, p < 1×10^-16^; Fig. 3B; Fig. S1B). In contrast, the gene body of highly expressed genes had substantially increased methylation (47.4% versus 14.9% in unexpressed genes; Wilcoxon rank-sum test: n_1_ = 333, n_2_ = 799, W = 70108, p < 1×10^-16^; Fig. 3A; Fig. S1A), and fewer open chromatin peaks (0.25 versus 0.94 open chromatin peaks per kilobase in unexpressed genes; Wilcoxon rank-sum test: n_1_ = 333, n_2_ = 842, W = 79825, p < 1×10^-16^; Fig.3B; Fig. S1B). The same pattern holds using a less strict cutoff for expression (75^th^ versus 25^th^ quantile; Fig S2) and when comparing expression deciles (Fig. S3).

**Figure 3.**
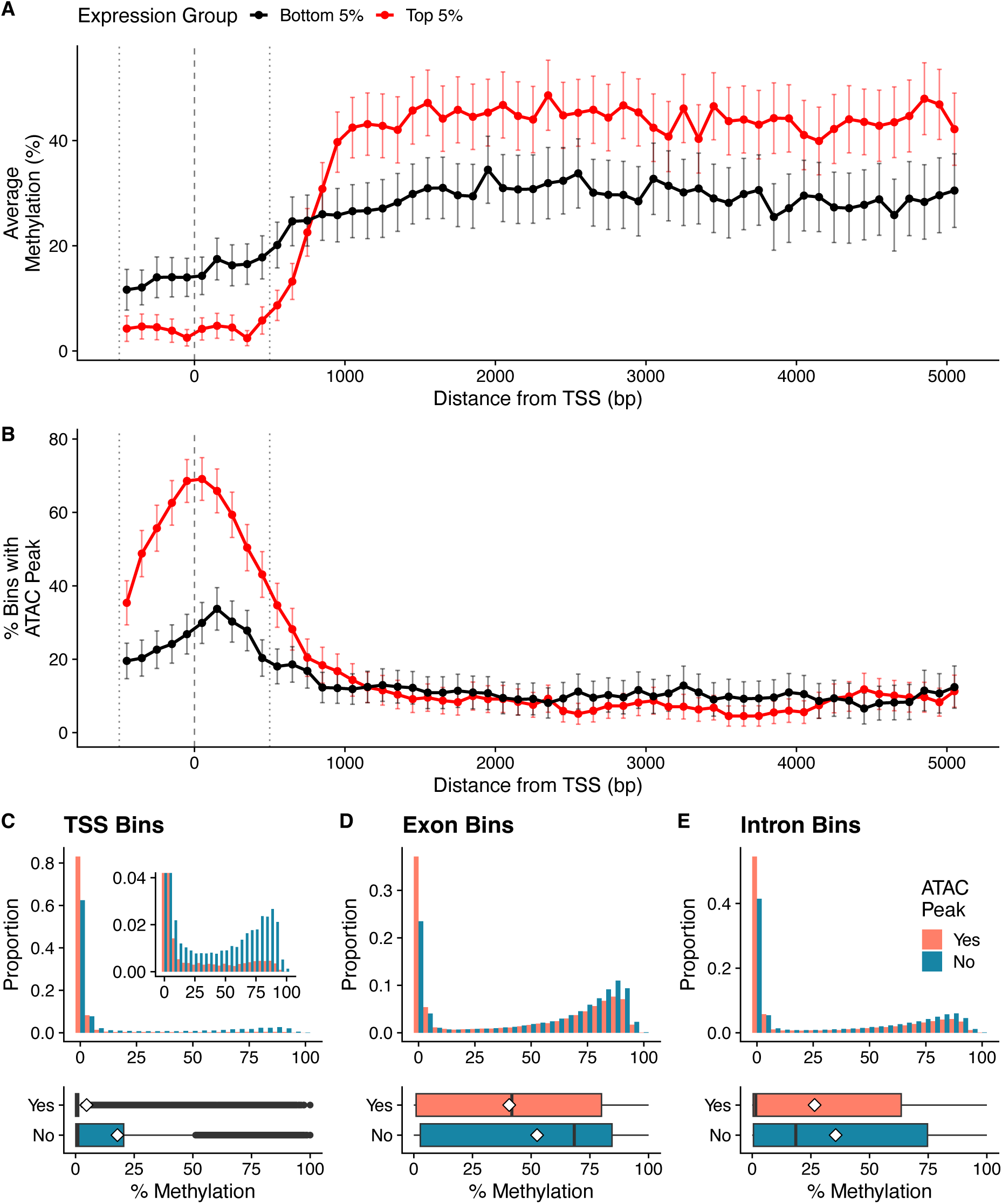
DNA methylation at the transcription start site (TSS) has a relationship with gene expression distinct from the rest of the gene body. (A) Highly expressed genes have reduced methylation at their TSS and increased gene body methylation, along with (B) a greater probability of having an open chromatin peak at their TSS. This relationship is modulated by the repression of open chromatin peaks by methylation at (C) the TSS, (D) exons, and (E) introns. (A-B) Points are averages for a 100 bp bin, across all genes within a given expression class, and error bars are the 95% CI. The vertical dashed line indicates the TSS, and the vertical dotted lines indicate the ‘TSS-adjacent’ region used in subsequent analyses. (C-E) Histograms show the percent methylation for 100 bp bins along all genes, dividing between bins containing and not containing an ATAC-seq peak. Boxplots show the same data, displaying median (center line), quartiles (box), and quartiles plus 1.5 times the interquartile range (whiskers), with diamonds indicating the average. (C) Exonic and intronic bins within 500 bp of the TSS. (D) Exonic bins more than 500 bp from the TSS. (E) Intronic bins more than 500 bp from the TSS.

Bins with greater methylation were consistently less likely to be in a peak of open chromatin. For example, bins adjacent to the TSS with more methylation had a greatly reduced chance of being in a region of open chromatin (Binomial-GLM, n = 47763: 𝛽 = −0.67, 95% CI = [−0.69, −0.64]; Likelihood ratio test: 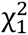 = 3503, p < 1×10^-16^; Fig. 3C). Greater methylation is also associated with less open chromatin in exon (Binomial-GLM, n = 78233: 𝛽 = −0.32 [−0.33, −0.30]; Likelihood ratio test: 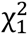 = 1297, p < 1×10^-16^; Fig. 3D), and intron (Binomial-GLM, n = 714934: 𝛽 = −0.26 [−0.27, −0.25]; Likelihood ratio test: 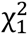 = 4322, p < 1×10^-16^; Fig. 3E) gene body bins, although the relationship is weaker in the gene body than at the TSS for both contexts. In summary, in *T. cristinae* highly expressed genes are typified by a TSS with low methylation and a peak of open chromatin, and a gene body with high methylation and few open chromatin peaks. We further examine how this pattern varies along the gene body, and how this is affected by gene length in the supplementary material (Fig. S4).

### Variation in methylation and expression between chromatin compartments

After establishing the epigenetic features which are most relevant to gene expression (open chromatin peaks and methylation at the TSS, and methylation on the gene body), we examined the covariation in these epigenetic features among genes and how they relate to chromatin compartments *i.e.*, regions of euchromatin and heterochromatin (Fig. 4). Note that while we discuss methylation adjacent to the TSS, this was highly correlated with both methylation at the promoter (r = 0.96, 95% CI = [0.96, 0.96]; t_6221_ = 272, p < 1×10^-16^) and at the first exon (r = 0.9, [0.89, 0.91]; t_1665_ = 86, p < 1×10^-16^). Despite a modest positive correlation between methylation around the TSS and gene body methylation (r = 0.37, [0.35, 0.39]; t_5918_ = 31, p < 1×10^-16^), they had opposite relationships with gene expression (r = -0.20, [-0.22, -0.17], t_5229_ = 14, p < 1×10^-16^, and r = 0.06, [0.03, 0.08], t_5379_ = 4.1, p = 4×10^-5^ respectively; Steiger’s test for differences between dependent correlation coefficient: t_5164_ = 16.7, p < 1×10^-16^; Fig. 5A-C). However, the difference between TSS methylation and gene body methylation did not predict expression better than TSS methylation alone (Steiger’s test: t_5164_ = 0.14, p = 0.89). This corroborates the findings from the previous section; wherein highly expressed genes are impoverished for methylation around the TSS and enriched for gene body methylation. Yet, it also suggests that when a gene is not being expressed, methylation at its promoter and gene body are typically in the same methylated/unmethylated state. Furthermore, chromatin openness at the TSS is more important for determining gene expression than the chromatin compartment a gene is in, with an open peak at the TSS having a stronger correlation with gene expression (r = 0.23, [0.2, 0.26], t_5230_ = 13.7, p < 1×10^-16^) compared to chromatin compartment score (r = 0.09, [0.07, 0.12], t_5448_ = 6.8, p = 1×10^-11^; Steiger’s test: t_5229_ = 8.7, p < 1×10^-16^; Fig. 5A, D, E). All correlations are qualitatively similar when using Spearman rank correlation (Fig. S5).

**Figure 4.**
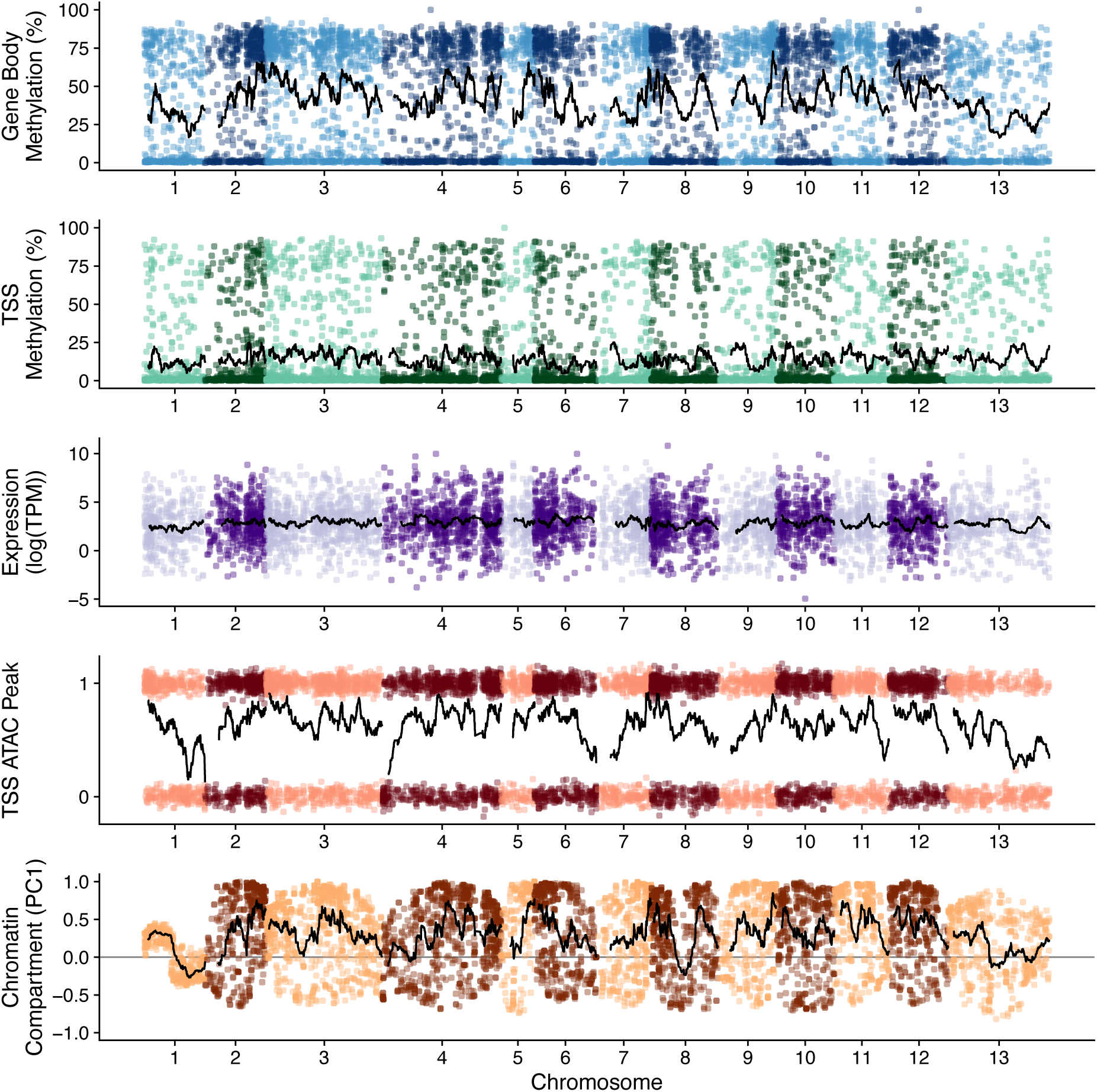
Patterns of epigenetic marks and gene expression for all analysed genes across the *T. cristinae* genome. Each point is a gene, with its x-position at the gene’s midpoint. Black lines are a moving average over 50 genes. ATAC peak presence is indicated by 1 and absence by zero, with points vertically jittered for visibility. Genes on alternating chromosomes are differently coloured for visibility.

**Figure 5.**
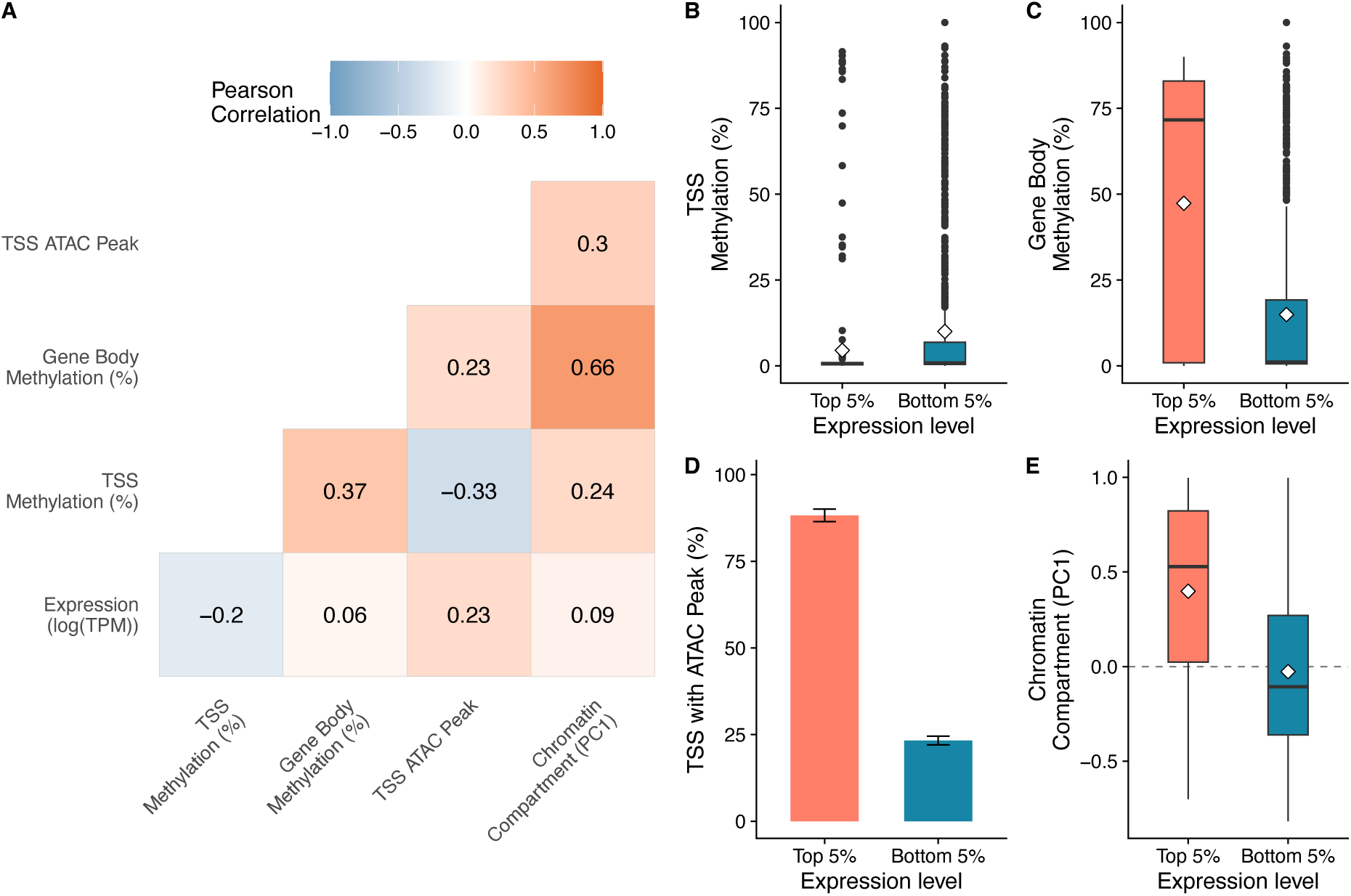
DNA methylation and open chromatin peaks correlate with both gene expression and chromatin compartments. (A) Pearson correlation among epigenetic features of genes and gene expression. A comparison of (B) transcription start site (TSS) methylation, (C) gene body methylation, (D) TSS openness and (E) chromatin compartment score, between highly expressed and unexpressed genes. (B, C, E) Boxplots display median (center line), quartiles (box), and quartiles plus 1.5 times the interquartile range (whiskers). The diamond indicates the mean. (D) Bar plot error bars are the 95% CI of the mean.

Genes occurring in regions of putative euchromatin were more likely to have open chromatin at their TSS than those in putative heterochromatin (r = 0.3, [0.27, 0.32], t_6369_ = 21, p < 1×10^-16^), as expected for genes in more active chromatin. However, the most striking trend was the large increase in gene body methylation in euchromatin compared to heterochromatin (48.7% versus 10.8% respectively; r = 0.66, [0.64, 0.67], t_6178_ = 69, p < 1×10^-16^), and an increase in TSS methylation in euchromatin (r = 0.24, [0.22, 0.27], t_6336_ = 20, p < 1×10^-16^) despite the negative association between TSS methylation and gene expression (Fig. 5A). These strong correlations were due to genes in heterochromatin compartments being nearly devoid of DNA methylation, whereas genes in regions of euchromatin had the bimodal pattern typical of methylation in invertebrates (Fig. 6A). Thus, the alternation between chromatin compartments nearly perfectly delineates between regions of high and low gene body methylation (Fig. 6B). Chromatin compartment also predicts non-gene body methylation better than gene body methylation, even though non-gene body methylation is not bimodally distributed (r = 0.73, [0.72, 0.73], t_11966_ = 115, p < 1×10^-16^; Steiger’s test: t_4720_ = 8.1, p = 7×10^-16^; Fig. S6).

**Figure 6.**
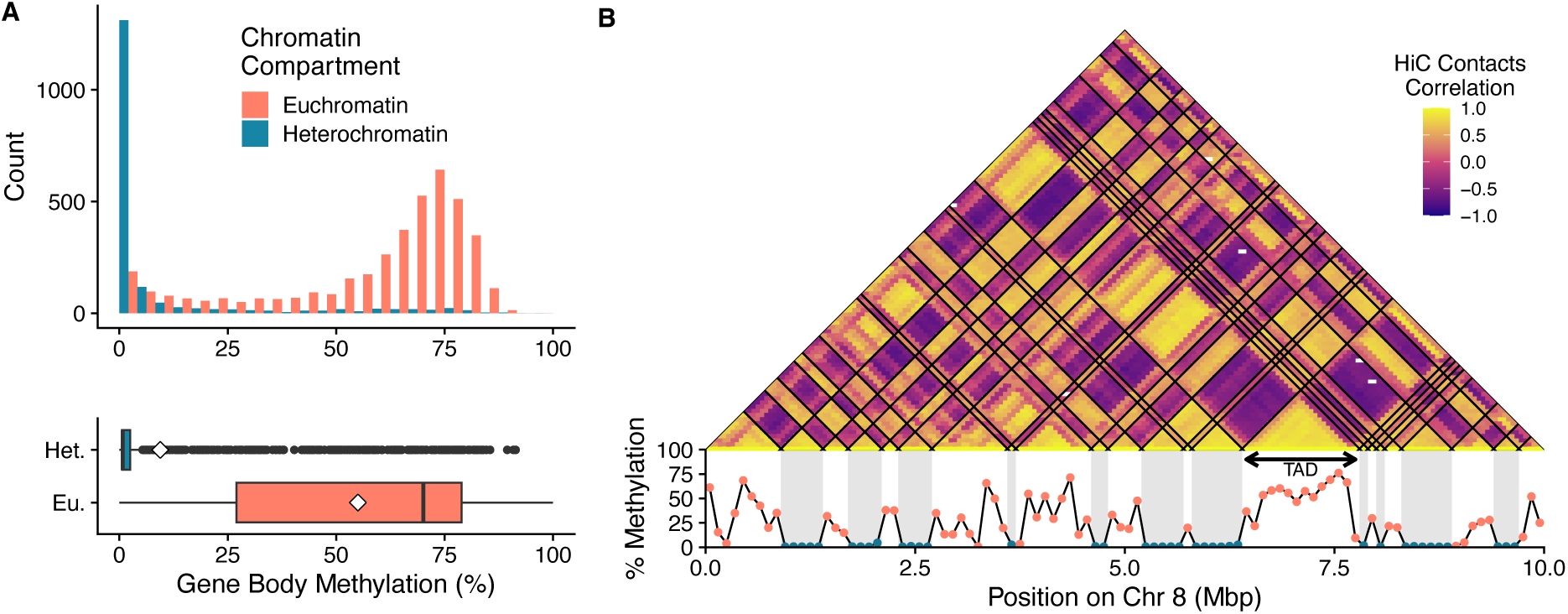
Heterochromatin compartments contain almost no DNA methylation. (A) A histogram and boxplot showing how the genome-wide bimodal pattern of DNA methylation divides into heterochromatin and euchromatin. White diamonds are the mean % methylation for each chromatin compartment. (B) An example of the pattern of chromatin contact autocorrelation from a 10 Mbp block on chromosome eight, overlayed with patterns of DNA methylation along the chromosome. A single topologically associated domain (TAD) was detected within this region and in annotated with a double-sided arrow. Black lines on the pyramid plot indicate annotated A/B compartment boundaries and shaded regions in the lower line plot are putative regions of heterochromatin.

The relationship between gene expression and TSS methylation is consistent between chromatin compartments whereas its relationship with gene body methylation differs. There is no significant interaction between TSS methylation and chromatin compartment when predicting gene expression (linear model, n = 5231: log(expression) ∼ TSS methylation + compartment score + TSS methylation × compartment score; TSS methylation, 𝛽 = -0.49, F_1, 5227_ = 190, p < 1×10^-16^; compartment score, 𝛽 = 0.32, F_1, 5227_ = 93, p < 1×10^-16^; TSS methylation × compartment score, F_1, 5227_ = 0.9, p = 0.35; Fig. S7). In contrast, there is no significant association between gene expression and gene body methylation after controlling for chromatin compartment, with a marginally significant interaction (linear model, n = 5381: log(expression) ∼ gene body methylation + compartment score + gene body methylation × compartment score; gene body methylation, F_1, 5377_ = 0.03, p = 0.86; compartment score, 𝛽 = 0.22, F_1, 5377_ = 27, p = 2×10^-7^; gene body methylation × compartment score, 𝛽 = 0.08, F_1, 5377_ = 3.74, p = 0.053; Fig. S7). These results suggest that chromatin compartments drive much of the association between gene body methylation and expression, but not TSS methylation and expression. This finding is corroborated by principal component analysis (PCA) of gene expression and epigenetic states together. Gene body methylation and chromatin compartments mostly load onto PC1, and expression and openness at the TSS mostly load onto PC2, whereas TSS methylation contributes substantially to both PCs (Fig. S8).

### Methylation at TAD boundaries

Due to the relatively low coverage of our Hi-C datasets, we detected a limited number of topologically associated domains (TADs) (452 TADs detected with a 10 kb sliding window). However, we examined methylation and chromatin openness at the detected TADs, as well as patterns at CTCF-binding motifs, to provide a preliminary look at whether methylation plays a role in meso-scale chromatin structure in *T. cristinae*. Methylation changed significantly more than expected by chance at TAD boundaries (ΔmCpG 24.3% at TAD boundaries versus 14.7% at random genomic positions; Wilcoxon rank-sum test: n_1_ = 901, n_2_ = 896, W = 486637, p = 5×10^-14^; Fig. S9A) but chromatin openness did not (openness was measured using ATAC-seq fragments per kilobase per million reads (FPKM); ΔFPKM at TAD boundaries 3.6 versus 3.4 at random genomic positions; Wilcoxon rank-sum test: n_1_ = 901, n_2_ = 887, W = 420356, p = 0.06; Fig. S9C). In contrast, at CTCF-binding motifs neither methylation nor chromatin openness shifted more than expected by chance (Wilcoxon rank-sum test: n_1_ = 1664, n_2_ = 1852, W = 1531219, p = 0.75 and n_1_ = 1655, n_2_ = 1828, W = 1564382, p = 0.08 respectively; Fig. S9B,D). But there is a clear peak of open chromatin at CTCF-binding motifs (2.9 FPKM at CTCF-binding motifs versus 0.02 FPKM at random genomic positions; Wilcoxon rank-sum test: n_1_ =1346, n_2_ = 1686, W = 1364367, p < 1×10^-16^; Fig S9D), and reduced methylation at CTCF-binding motifs that overlap a peak of open chromatin (8.9% methylation at CTCF-binding versus 11.4% methylation at random genomic positions; Wilcoxon rank-sum test: n_1_ = 222, n_2_ = 1434, W = 182334, p = 5×10^-4^; Fig. S10). These results lend further credence to a role of methylation in structuring chromatin compartments, potentially through interaction with CTCF-binding factors as reported in mammals^26,38,56^.

### Comparative analysis of Apis mellifera

To assess the generality of our findings in *T. cristinae*, we performed the same analyses in *A. mellifera*, using publicly available data. *Apis mellifera* is the most widely used insect model for studies of DNA methylation, having been used to study the role of DNA methylation in learning, epigenetic inheritance^57,58^, caste differentiation^59^, and many other life history traits. Furthermore, there are publicly available RNA-seq^60,61^, BS-seq^62^, ATAC-seq^61^, and Hi-C^60^ data for this species, allowing us to directly compare to our analysis in *T. cristinae*. Importantly, the pattern of DNA methylation in *A. mellifera* differs from that of *T. cristinae*, with DNA methylation in *A. mellifera* occurring nearly exclusively in exons^63^. This pattern is likely representative of most holometabolous insects, which have highly reduced DNA methylation^45,64^.

In *A. mellifera*, TSS methylation did not correlate with gene expression, but gene body methylation did. Both highly expressed and unexpressed genes had almost no TSS methylation (1.3% versus 0.5% methylation, respectively; Wilcoxon rank-sum test: n_1_ = 253, n_2_ = 276, W = 31266, p = 0.04; Fig. 7A; Figs. S11A, S12A). In contrast, highly expressed genes had elevated gene body methylation, with the maximum around 1000 bp downstream of the TSS, whereas unexpressed genes largely lacked gene body methylation (29.2% versus 4.2% methylation, respectively; Wilcoxon rank-sum test: n_1_ = 334, n_2_ = 234, W = 12050.5, p < 1×10^-16^; Fig. 7A; Figs. S11A, S12A). As in *T. cristinae*, highly expressed genes were more likely to have a peak of open chromatin at their TSS than unexpressed genes (38.8% versus 20.3% of TSSs with an open chromatin peak, respectively; Wilcoxon rank-sum test: n_1_ = 286, n_2_ = 255, W = 29703, p = 2×10^-6^; Figs. S11B, S12B). Also as in *T. cristinae*, greater methylation was associated with a lower probability of an open chromatin peak at the TSS (Binomial-GLM, n = 31722: 𝛽 = −0.30 [−0.36, −0.24]; Likelihood ratio test: 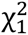 = 220, p < 1×10^-16^; Fig. S11C), exons (Binomial-GLM, n = 104086: 𝛽 = −0.69 [−0.71, −0.66]; Likelihood ratio test: 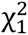 = 3833, p < 1×10^-16^; Fig. S11D), and introns (Binomial-GLM, n = 329403: 𝛽 = −0.21 [−0.24, −0.18]; Likelihood ratio test: 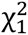 = 424, p < 1×10^-16^; Fig. S11E). However, methylation at the TSS and in introns was extremely low. The same pattern holds using a less strict cutoff for expression (75^th^ vs 25^th^ quantile; Fig. S13) and when comparing expression deciles (Fig. S14). As with *T. cristinae*, we further explore changes in methylation along the gene body and its relationship with gene length in *A. mellifera* in the SI (Fig. S15). These findings suggest that in *A. mellifera* gene body methylation rather than TSS methylation is important in the regulation of gene expression.

**Figure 7.**
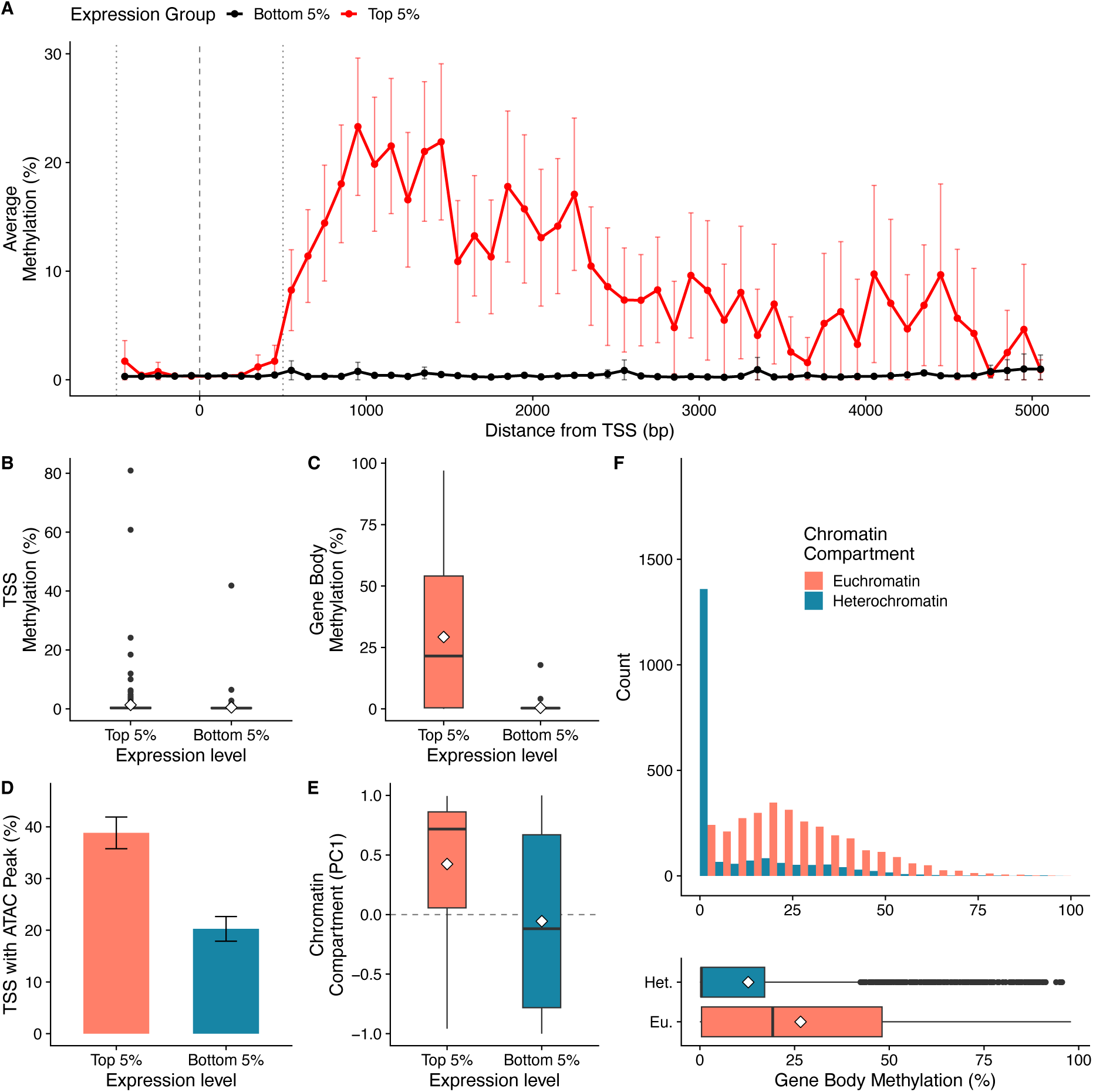
DNA methylation in *Apis mellifera* has a different relationship with gene expression but shows the same relationship with chromatin compartments as in *Timema cristinae*. (A) Highly expressed genes have increased gene body methylation but no difference at the transcription start site (TSS). Points are averages for a 100 bp bin, across all genes within a given expression class, and error bars are the 95% CI. The vertical dashed line indicates the TSS, and the vertical dotted lines indicate the ‘TSS-adjacent’ region used in subsequent analyses. A comparison of (B) TSS methylation, (C) gene body methylation, (D) TSS openness and (E) chromatin compartment score, between highly expressed and unexpressed genes. (F) A histogram and boxplot showing how the genome-wide bimodal pattern of DNA methylation divides into heterochromatin and euchromatin. White diamonds are the mean % methylation for each chromatin compartment. All boxplots display median (center line), quartiles (box), and quartiles plus 1.5 times the interquartile range (whiskers). Diamonds indicates the mean. Error bars on bar plots are the 95% CIs.

The above findings are corroborated by a weaker correlation between TSS methylation and expression (r = 0.04 [0.01,0.06]; t_5036_ = 2.6, p = 0.009), than between gene body methylation and expression (r = 0.35 [0.33,0.37]; t_6587_ = 30, p < 1×10^-16^; Steiger’s test: t_4911_ = 19, p < 1×10^-16^; Fig. 7B-C; Fig. S16A). Interestingly, gene expression was also more strongly correlated with chromatin compartment score (r = 0.23 [0.20, 0.25]; t_6777_ = 19, p < 1×10^-16^) than having an open peak at the TSS (r = 0.12 [0.09,0.15]; t_5049_ = 7.5, p = 7×10^-14^; Steiger’s test: t_5048_ = 5.3, p = 1×10^-7^; Fig. 7D-E; Fig. S17A). Results remain qualitatively the same using Spearman’s correlations (Fig. S16B).

In *A. mellifera*, DNA methylation is elevated in euchromatin compartments, but chromatin compartments do not explain the association between gene body methylation and gene expression. There was both greater TSS methylation (r = 0.09 [0.06, 0.12]; t_5161_ = 6.6, p = 6×10^-11^) and gene body methylation (r = 0.26 [0.23, 0.28]; t_6654_ = 22, p < 1×10^-16^; Fig. 7F) in euchromatin compartments than in heterochromatin compartments. However, TSS methylation had a weak relationship with gene expression regardless of chromatin compartment (linear model, n = 5038: log(expression) ∼ TSS methylation + compartment score + TSS methylation × compartment score; TSS methylation, 𝛽 = 0.07 [0.01, 0.13], F_1, 5034_ = 4.6, p = 0.03; compartment score, 𝛽 = 0.46 [0.40, 0.52], F_1, 5034_ = 241, p < 1×10^-16^; gene body methylation × compartment score, 𝛽 = -0.1 [-0.18, -0.03], F_1, 5034_ = 7.5, p = 0.006; Fig. S17). Whereas, gene body methylation maintained a strong positive association with gene expression regardless of chromatin compartment (linear model, n = 6589: log(expression) ∼ gene body methylation + compartment score + gene body methylation × compartment score; gene body methylation, 𝛽 = 0.68 [0.64, 0.73], F_1, 6585_ = 760, p < 1×10^-16^; compartment score, 𝛽 = 0.27 [0.22, 0.32], F_1, 6585_ = 111, p < 1×10^-16^; gene body methylation × compartment score, 𝛽 = - 0.22 [-0.27, -0.17], F_1, 6585_ = 68, p < 1×10^-16^; Fig. S17).

This comparison of gene expression, DNA methylation and chromatin state between *T. cristinae* and *A. mellifera* suggest that some aspects of DNA methylation are consistent among insects while others are not. The presence of TSS methylation, and the function of gene body methylation differ between the two species compared, possibly representing a shift in the function of gene body methylation in Holometabola. In contrast, DNA methylation is elevated in euchromatin compartments in both species, potentially representing a conserved pattern.

### The ancestral function of DNA methylation in Animalia

As a preliminary investigation of the phylogenetic history of TSS and euchromatin methylation, we compared our findings in *T. cristinae* and *A. mellifera* to findings from studies of DNA methylation in other invertebrates and some vertebrates (Table S1). We defined the presence of TSS methylation as a negative association between gene expression and DNA methylation at the promoter or regions flanking the TSS, and we defined euchromatin methylation as elevated methylation in euchromatin compartments relative to euchromatin compartments. This analysis suggests that functional TSS methylation and ‘mosaic’ euchromatin methylation are the ancestral states of DNA methylation in animals (Fig. 8; Table S1). The presence of TSS methylation in *T. cristinae*, along with many arthropods outside of Holometabola, and all non-arthropod animals studied so far suggest that the common ancestor of animals likely had functional TSS methylation (posterior probability 87%; Fig. 8A). This also implies that within Arthropoda, TSS methylation has been lost and gained at least twice. On the other hand, DNA methylation is associated with active euchromatin in all invertebrates, making it likely (posterior probability 82%) that the common ancestor of animals had DNA methylation limited to regions of euchromatin (Fig. 8B). These findings also indicate uncertainly about whether the common ancestor of eukaryotes had predominantly euchromatin or heterochromatin methylation, but more complete phylogenetic sampling is required to draw any definitive conclusions.

**Figure 8.**
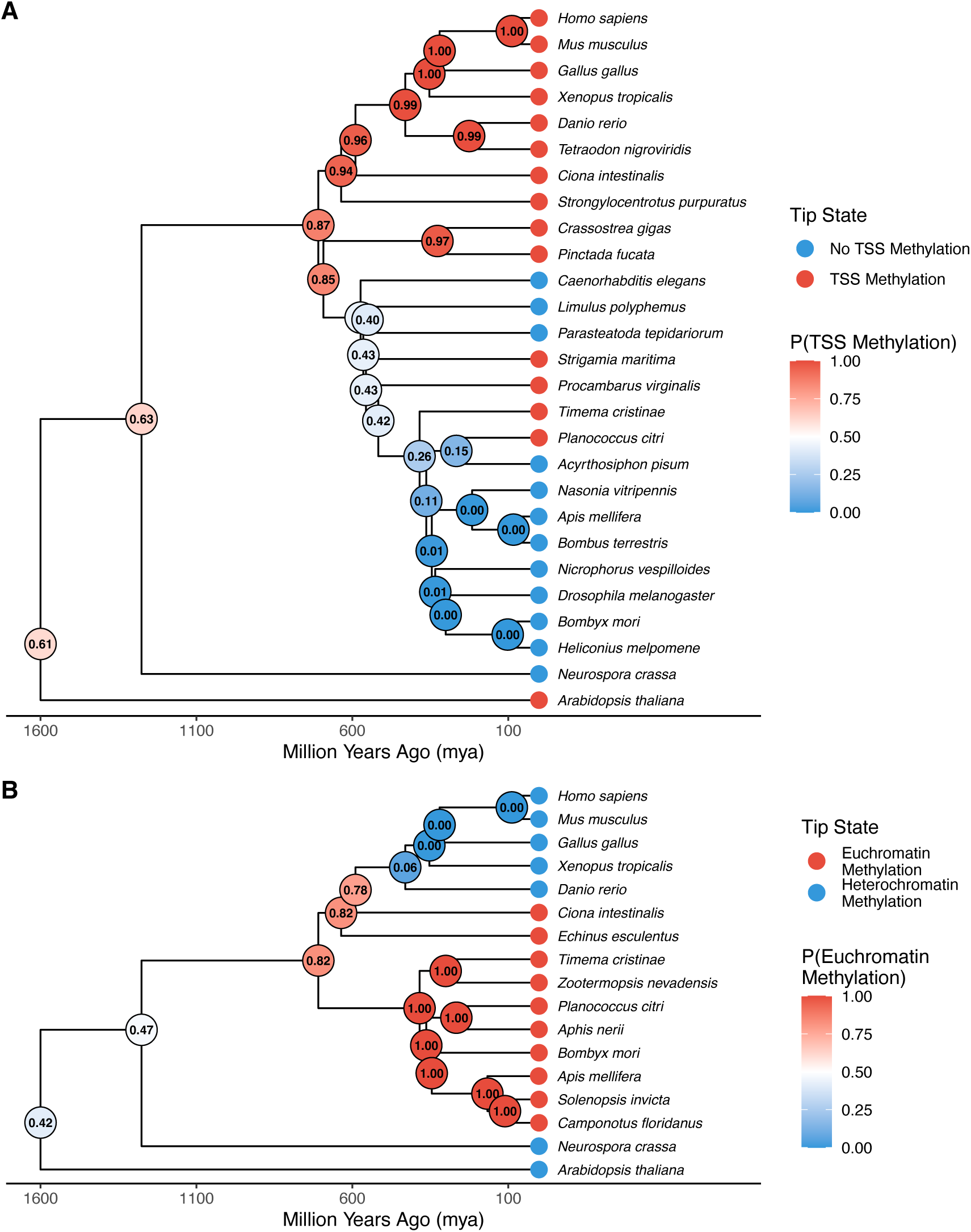
Transcription start site (TSS) methylation and euchromatin methylation are common among invertebrates. Ancestral state reconstruction for (A) transcription start site (TSS) methylation, and (B) euchromatin methylation, across Eukaryota. Nodes are labeled with posterior probability based on 1000 simulations of stochastic character mapping with equal rates (ER). (A) The posterior probability for the node between *C. elegans* and Arthropoda is 0.43.

## Discussion

### Summary

Here, we have integrated sequencing of genome-wide DNA methylation, chromatin openness, 3D topology, and gene expression in a stick insect. Using these data, we provide a detailed analysis of the mechanisms through which DNA methylation influences gene expression in a stick insect and make inferences about invertebrates more broadly. Our findings support the hypothesis that in *T. cristinae*, DNA methylation is involved in gene expression at the TSS, rather than the gene body (Hypothesis 1; Fig. 2A). We showed that methylation at the transcription start site is negatively associated with gene expression, whereas gene body methylation is positively associated with gene expression, as in many non-insect invertebrates^6,11^. However, the positive correlation between gene body methylation and expression, appears to largely be driven by gene body methylation targeting regions of active chromatin. Our findings also suggest that in *T. cristinae*, DNA methylation is associated with both open and closed chromatin, depending on the scale (Hypothesis 2; Fig. 2B). At the scale of chromatin compartments, DNA methylation occurs in more open and active regions of euchromatin. However, within these active compartments, DNA methylation is associated with less accessible regions of DNA, at both the TSS and gene body, suggesting that methylation influences gene expression through the stabilization of nucleosome complexes^65^. Lastly, comparative analysis with *A. mellifera* suggests that while the function of TSS methylation has changed among insect species DNA methylation has maintained the same relationship with chromatin. These findings are consistent with studies of other invertebrate species using electrophoresis^42,43^, or comparing methylation to histone modifications^66–68^. Together these observation suggest that the relationship between heterochromatin and DNA methylation in invertebrates is opposite of that of vertebrates^69^. Thus, TSS methylation may be more common among invertebrates than previously appreciated but not ubiquitous, and chromatin compartments are potentially an essential feature structuring invertebrate DNA methylation across the genome.

### TSS methylation in invertebrates

TSS methylation is potentially widespread among invertebrates. The low levels of DNA methylation in invertebrates, and the heterogeneous relationship between TSS methylation and expression has led to a general consensus that there is little to no TSS methylation in invertebrates^11,14^. However, there are observations of negative correlations between promoter/TSS methylation and gene expression from Tunicata (*Ciona intestinalis*^6^), Mollusca (*Crassostrea gigas*^54,70,71^ but see Olson and Roberts^72^ for contrasting evidence), Echinodermata (*Strongylocentrotus purpuratus*^73^, and species from three different classes of arthropods (*Planococcus cirtri*, *Strigamia maritima*, and *Procambarus virginalis*^13,14^). Furthermore, Gatzmann et al.^13^ and Bogan et al.^73^ also compared peaks of open chromatin to DNA methylation and expression using ATAC-seq, both finding the same negative association between DNA methylation and open chromatin at the TSS as we do here. These findings in addition to ours here suggest that TSS methylation may be common among invertebrates and involved in the regulation of chromatin, but lost in some groups such as Holometabola.

Our findings suggest a possible explanation for inconsistent correlations between TSS methylation and gene expression among invertebrates. As there is reduced DNA methylation in heterochromatin compartments, unexpressed genes in these compartments are more likely to have low TSS methylation. Whereas unexpressed genes in euchromatin compartments have elevated TSS methylation, typical of the repression of expression by DNA methylation. Since euchromatin makes up 73% of the *T. cristinae* genome, the relationship between methylation and expression in these compartments drives the overall trend and we see a positive correlation. Whereas, if other invertebrates have substantially more heterochromatin, the overall association would be substantially weaker or even reversed. Therefore, future investigation into the relationship between DNA methylation and gene expression in invertebrates should take chromatin compartments into consideration. Thankfully, chromatin compartments can be estimated from DNA methylation. For example, using a threshold of 5% methylation to designate chromatin compartments assigns 86% of 100 kb bins to the correct chromatin compartment in *T. cristinae* (Fig. S18). Furthermore, using the correlation in DNA methylation among biological replicates can predict chromatin compartments even more accurately^74^.

Methylated TSSs in invertebrates have overall low levels of methylation, suggesting that TSS methylation may not be functional in the repression of gene transcription in this clade. However, it is unknown whether low levels of TSS methylation in invertebrates are due to TSS methylation being cell-type-specific or occurring stochastically among all cells at a low rate. If the former is true, TSS methylation would be more likely to have a functional role in gene expression. Tissue-specific patterns of DNA methylation are supported by the coincidence of changes in life history stage and DNA methylation among many invertebrates^70,75–77^. Furthermore, the strength of correlation between invertebrates TSS methylation and gene expression is of the same order of magnitude as similar analyses in humans and other vertebrates, with Pearson and Spearman correlations ranging from around zero to -0.45^78–80^. However, there are also a growing number of observations that invertebrate DNA methylation is relatively invariable between tissue types^13,72,81,82^), and there is evidence against differences in in DNA methylation among insect castes^81,83^. However, the analyses cited above focus on overall patterns of DNA methylation, which are largely driven by gene body methylation (the predominant form of DNA methylation in invertebrates), whereas TSS methylation may show a distinct tissue-specific pattern. Interestingly enough, invertebrate genes with gene body methylation are more likely to retain housekeeping function in vertebrate orthologs, whereas genes with TSS methylation are more likely to be tissue-specific in vertebrate orthologs^6^. This suggests that TSS methylation should be considered separately from gene body methylation, when assessing the function and evolution of invertebrate DNA methylation (Fig. 3).

### Chromatin compartment methylation

The ‘mosaic’ pattern of DNA methylation in invertebrates is likely due to chromatin compartments. While we are the first (to our knowledge) to explicitly compare chromatin compartments to DNA methylation data in an invertebrate, it is well established that methylation occurs in a mosaic pattern in invertebrates^9,11^. For example, Glastad et al.^76^ described the pattern of methylation in the termite *Zootermopsis nevadensis,* as “clusters of methylated genes [in] regions up to hundreds of kilobases in size”. This pattern exactly corresponds with our observations of highly methylated regions being nearly exclusive to gene-rich compartments of euchromatin on the order of hundreds of kilobases. Furthermore, Bird et al.^84^ described similar 100 kb scale blocks in DNA methylation of the sea urchin *Echinus esculentus*. These observations suggest that compartmentalization is a major driver of patterns of DNA methylation across invertebrates but further analysis among diverse taxa is required to verify this claim.

### Discerning causality

Our results do not directly inform us about the causal relationship between DNA methylation, gene expression, and chromatin compartment formation. While we observe an association between TSS methylation and expression, as well as methylation and chromatin compartments, this only suggests that methylation could cause either. As has been previously discussed and debated^85–87^, gene body methylation may be a response to gene expression and changes in chromatin state, rather than a precursor. This is supported by evidence that changes in expression occur prior to changes in DNA methylation, during X chromosome inactivation and chromatin remodeling^34,88^. Furthermore, alteration of gene body methylation does not affect gene expression in *A. thaliana*^89^ but findings from methylation knockdown experiments in invertebrates are mixed^90,91^. As targeted epigenetic editing tools become more accessible for use in non-model species, it will become possible to test the functional role of DNA methylation in a wider taxonomic range of invertebrates^92,93^.

This debate also extends to the role of DNA methylation in heterochromatin compartment formation. Despite the well-established role of DNA methylation in heterochromatin regulation^94,95^, evidence suggests that DNA methylation is implicated in the stabilization of heterochromatin, rather than the initiation of its formation^96^. From this we infer that the lack of methylation in the heterochromatin compartments of *T. cristinae* is a result of the formation of stable heterochromatin, rather than a precursor to its formation. While such a relationship can be tested causally by artificially modifying TAD structure^97,98^, current methods require the use of transgenic lines and are limited to model species.

## Conclusion

We found that DNA methylation at the transcription start site of genes in the stick insect *T. cristinae* is negatively associated with gene expression and chromatin accessibility. Despite this, DNA methylation was nearly absent from heterochromatin compartments. This suggests that invertebrate DNA methylation has retained functions similar to those of vertebrate and plant methylation, with respect to gene expression. However, invertebrate DNA methylation is highly divergent or non-functional with respect to heterochromatin formation. Further studies of phylogenetically diverse non-model invertebrates will further elucidate the exact phylogenetic history of the relationship between DNA methylation and gene expression. Furthermore, the growing accessibility of Hi-C data from chromosome-level genome assemblies in non-model organisms can be used to gain deeper insights into the molecular regulation of gene expression and how chromatin structure and the epigenetic landscape shape evolution. By examining how DNA methylation has evolved to fulfill a wide variety of functions in a stick insect, we better understand the evolutionary history of gene expression, its regulation, and how to better predict evolution in the future.

## Materials and Methods

All datasets were mapped to haplotype one of the Hwy154 striped reference described in Gompert et al.^53^, using only chromosome-level scaffolds (genome ID: CEN4280; NCBI bioproject number: PRJNA1208028). Gene annotation was done using BRAKER3 with the exact same protocol as Gompert et al., (2025), except that we used the UTR=on setting for more precise TSS annotation^99^. After filtering, there were 6,663 high-confidence non-overlapping gene annotations. All code will be made available at https://github.com/planidin/Tcris_chromatin and data will be uploaded to Zenodo.

### RNA-seq

Raw paired-end RNA-seq data were generated from 18 adult female *Timema cristinae* as part of a previous study^52^ (average 7.08M reads; Table S2). Raw reads were trimmed for adapter content (see code for adapter sequences), quality (-q 20) and length (-m 35) using *Cutadapt* 1.16^100^. Then again with *Trimmomatic* 0.39^101^, trimming for adapter content (ILLUMINACLIP: Illumina_TruSeq.fa:2:30:8:1:true), length (CROP:150, MINLEN:35) and quality (LEADING:20, TRAILING:20, SLIDINGWINDOW:4:20, AVGQUAL:20). We then mapped reads using *STAR* 2.7.11b^102^ in basic two-pass mode, mapping all reads in the first step, filtering out mappings with greater than 5% mismatch to mapped read length (--outFilterMismatchNoverLmax 0.05) and more than 20 mismatches (--outFilterMismatchNmax 20). We then filtered out multimapping reads (-q 20) and unpaired reads (-f 3) using *samtools* 1.16^103^, resulting in an average of 3.27M reads per individual (Table S2). Finally, gene expression was tabulated using *featureCounts* 1.5.3^104^ in paired-end mode (-p), counting fragments where both reads aligned to the same chromosome (-B, -C). Fragment counts normalized to transcripts per million (TPM) using a custom bash script were then used for downstream analysis.

### BS-seq and Methyl-seq

We use paired-end BS-seq from 24 adult female *T. cristinae* from a previous study^52^ (average 40.59M reads; Table S3) and newly generated paired-end Methyl-seq from 10 additional adult females (average 107.42M reads; Table S3). Raw reads were first trimmed using *TrimGalore* 0.6.6^100^, removing adapter content, low-quality bases (--quality 20), and overly short reads (--length 36). Reads were mapped using *Bismark* 0.24.1^105^ in paired-end mode, with twenty consecutive extension attempts (-D 20), three reseed attempts (-R 3), and otherwise default settings. After mapping there was an average of 30.14M reads per individual (Table S3). Methylation percentages were then calculated with *bismark_methylation_extractor* ignoring the first seven and last two positions of both reads (--ignore 7, --ignore_r2 7, -- ignore_3prime 2, --ignore_3prime_r2 2) for BS-seq samples and ignoring the first three and last position of both reads for Methyl-seq samples, due to bias in methylation levels at these positions (Fig. S19). This resulted in a final average of 164.7M reads at CpG sites per individual (Table S3). As methylation levels at CHG and CHH motifs match error rate expected by bisulfite conversion 0.2%^52^, only CpG methylation was used for subsequent analysis.

We calculated the average methylation at each CpG site using a weighted average, normalizing for coverage. First, cytosine (C) and thymine (T) counts were normalized to counts per million reads (CPM), to account for differences in coverage among individuals. Then the normalized C and T counts were summed across individuals, and the average percent methylation was tabulated from this sum. We then removed CpG sites with coverage less than or greater than 2 standard deviations from the mean, as these were under and over methylated respectively (Fig. S20). This method allows for many more CpG sites to be included in analysis (50,099,801 sites) compared to taking the average across individuals for CpG sites with at least five reads for all individuals (32,535 sites) and gives nearly the same estimate of methylation (r = 0.998; Fig. S21).

We used a similar weighted average technique to calculate the average methylation of genomic features e.g., gene bodies. The C and T counts of all CpG sites within a genomic feature were summed, then we calculated the overall percent methylation from this sum. Thus, CpG sites with a more accurate estimation of percent methylation (higher coverage) are more heavily weighted in the average. Percent methylation was consistent between BS-seq and Methyl-seq samples at CpG sites (r = 0.92; Figure S22) and highly consistent for gene body averages (r = 0.99; Figure S23). See code for details and implementation.

### ATAC-seq

Flash-frozen tissue samples from 10 adult female *T. cristinae* were shipped to Active Motif for ATAC-seq processing in two batches. Upon receipt, tissues were submerged in cold PBS and minced using a sterile razor blade. The minced tissue was homogenized in ATAC Lysis Buffer with a Dounce homogenizer. The resulting lysate was passed through a 40 µm mesh strainer to isolate nuclei. Nuclei were counted, and 100,000 nuclei were subjected to tagmentation using the enzyme and buffer supplied in the ATAC-Seq Kit (Active Motif). Tagmented DNA was purified using a DNA extraction method, followed by PCR amplification. The resulting libraries were purified with SPRI beads, quantified, and sequenced on the Illumina platform using paired-end 50 bp (PE50) reads (average 41.33M reads per sample; Table S4). Prior to sequencing, fragments were size selected to be < 150 bp, maximizing the sensitivity of detecting open chromatin peaks relative to coverage. However, this means that our ATAC-seq data does not capture information about nucleosome positioning.

Reads were first trimmed using *TrimGalore* with the same settings as described above. After trimming, forward and reverse reads were mapped separately using *aln* in *BWA* 0.7.17^106^, then merged using *sampe*. Following mapping, duplicate reads were removed using *Picard* 3.3.0^107^ *MarkDuplicates* and *samtools view* (-F 1024), then filtered for quality (-q 20), proper pairing (-f 3) using *samtools fixmate* and *samtools view*. After filtering there was an average of 28.86M reads per individual (Table S4). We then shifted forward and reverse reads by 4 and - 5 bp respectively, to account for the Tn5 insertion bias, using a custom python script. Open chromatin peaks were called using *MACS* 2.2.7^108^ *callpeak* in paired-end mode. Individual peaks were then merged into a set of consensus peaks using *BEDTools* 2.28.0^109^ *merge*, resulting in 187,826 open chromatin peaks (Table S4). Finally, fragment counts per peak and TSS enrichment were calculated using *samtools*, *BEDTools* and a custom bash script. This resulted in an average fragments-in-peaks (FRiP) score of 0.17 and TSS enrichment of 5.43 (Table S4; Fig. S24). To calculate the average fragment counts of genomic features for downstream analysis, fragment counts for each individual were first normalized to fragments per million fragments (FPM), then averaged across individuals.

Regions of open chromatin are consistent between sequencing batches and *T. cristinae* ecotypes. To ensure the consistency of open chromatin regions among individuals, we performed differential accessibility analysis. To do so, we downsampled the reads of all individuals to have the same coverage as the lowest coverage individual. Then, we called a new set of consensus peaks, resulting in 88,958 open chromatin peaks. Fragment counts were consistent between batches (r=0.84; Fig. S25). Furthermore, we used *DESeq2*^110^ to compare sequencing batches and host-plant ecotypes. We found that 3.3% of peaks were differentially accessible between sequencing batches (2,961 peaks; Fig. S26A-C), and one differentially accessible peak between host-plant ecotypes (Fig. S26D-F).

### Hi-C

Two Omni-C (182.21M and 146.56M reads) and one Hi-C (260.57M reads) dataset were generated as part of previously published *de novo* genome assemblies^53,111^ (Table S5). We first trimmed reads using *TrimGalore* with the same settings described above. Following trimming, we mapped reads and generated contact matrices using *Juicer* 1.5.6^112^, resulting in an average of 82.12M intra-chromosomal contacts per individual. We then calculated AB-compartment scores (PC1 of the correlation matrix of contacts) for each sample in 100 kb bins, using *JuicerTools eigenvector* with Knight-Ruiz normalization (KR). Bins were determined to be putative-euchromatin or putative-heterochromatin by correlating AB-compartment score with gene density. Compartment assignment by gene density has a high concordance with compartment assignment by gene expression, open chromatin peak/fragment count, gene body/CpG methylation, and CpG count (97.4% to 100% matching bins among methods). Furthermore, there was little variation in contact patterns and compartment assignment among samples (91% to 98% matching bins among individuals; for an example see Fig. S27). So, we used the average normalized AB-compartment scores for downstream analysis. AB-compartment assignment was done using a custom R script. See code for implementation.

### Within-gene analysis

We examined the pattern of epigenetic marks along genes, summarizing by 100 bp bins. We generated 100 bp bins along gene bodies using *BEDTools*, including the 500 bp upstream of the TSS. We chose 500 bp as the cutoff for upstream bins because this captures the region around the TSS which is consistently elevated for ATAC-seq fragments (open chromatin; Fig. S24), and this cutoff has been used in other studies^6^. Then, for each bin we calculated the average CpG methylation using the weighted average method described above and determined whether the bin overlapped with an ATAC-seq peak. All datasets were then summarized into a single table using a custom bash script and analysis performed in *R* 4.5.0. Methylation and ATAC-seq peaks within bins were compared using a binomial *glm* with scaled percent methylation as the sole predictor. This was done separately for bins within 500 bp of the TSS (regardless of exon or intron content), and exons and introns, more than 500 bp from the TSS.

### Among-gene analysis

Following within-gene analysis, we calculated the number of open chromatin peaks per kilobase and average percent CpG methylation within 500 bp of the TSS, and for the rest of the gene body, using *BEDTools*. All datasets were then summarized into a single table using a custom bash script and statistical analysis and plotting was performed in *R* 4.5.0. As 95.8% of TSS-adjacent regions contained either one or zero open chromatin peaks, we converted chromatin peaks per kilobase to a binary (none or at least one peak) for analysis. Differences in Pearson correlation were tested for statistical significance using Steiger’s test, implemented in the function *paired.r* from *psych* 2.5.3^113,114^. Principal component analysis was done with the built-in R function *prcomp*.

### TAD and CTCF-binding site analysis

We annotated topologically associated domains using the Hi-C contacts from the Hwy154 striped reference chromosomal assembly. We used *JuicerTools Arrowhead* with 10 kb windows (-r 10000), moving in 2 kb increments (-m 2000), with KR normalization (-k KR) and ignoring sparsity (--ignore_sparsity). We detected 452 TADs, ranging from 120 kb to 3.6Mb with a median size of 230 kb. We then summarized ATAC-seq fragments per kilobase per million reads (FPKM) and CpG methylation for 100 bp windows in the 50 kb upstream and downstream of each TAD boundary. This size range was chosen to capture as much of each TAD as possible, without overlapping. We then performed the same summarized for 452 random sites in the genome to compare to TAD boundaries.

We annotated putative CTCF-binding motifs in our reference genome using the position weight matrix (PWM) for CTCF binding in *Drosophila melanogaster* from Holohan et al.^115^. Motif scores for this PWM were calculated using *FIMO* in *MEME suite 5.5.7*^116^. ATAC-seq FPKM and CpG methylation were then summarized for 100 bp windows in the 50 kb surrounding each of 1,858 high-confidence CTCF-binding motifs (p < 10^-6^). We then generated the same summaries for 1,858 random sites in the genome to compare to CTCF-binding motifs.

### Comparative analysis of Apis mellifera

All *A. mellifera* sequencing datasets were from whole-body tissue samples of mature workers. We used RNA-seq data from Lowe et al.^61^ (n = 2; Table S7) and Jin et al.^60^ (n = 3; Table S7), BS-seq data from control individuals in Rasmussen et al.^62^ (n = 5; Table S8), ATAC-seq data from Lowe et al.^61^ (n = 2; Table S9), and Hi-C data from Jin et al.^11760^ (n = 1; Table S10). All datasets were mapped to the chromosomal-level scaffolds of the Amel_HAv3.1 reference genome (GCF_003254395.2), using the same quality control, and mapping pipeline as with *T. cristinae*. All mapped sequences were summarized using the same pipeline as with *T. cristinae*, except that BS-seq reads were trimmed for their first seven and last two positions (Fig. S28). There was also less bias in methylation at high coverage CpG sites in the *A. mellifera* dataset (Fig. S29). Furthermore, the ATAC-seq enrichment peak occurred downstream of the TSS, rather than upstream of the TSS as in *T. cristinae* (Fig. S30). Non-overlapping gene annotations from the Amel_HAv3.1 archive were used for downstream analyses. For accession numbers, and summarization of datasets, see Tables S7-10.

### Ancestral state reconstruction

We performed ancestral state reconstruction on two traits (i) the presence of TSS methylation which covaries with gene expression, and (ii) methylation in euchromatin compartments. Sources for phenotypic information for each species are given in Table S1. A chronogram was generated using https://timetree.org. Phylogenetic positions without data on timetree were approximated using another species in the same genus if possible: *Strongylocentrotus purpuratus* was estimated with *Strongylocentrotus intermedius*; *Ciona intestinalis* was estimated with *Ciona savignyi*; *Bombus terrestris* was estimated with *Bombus sp.*; *Heliconius melpomene* was estimated with *Heliconius ethilla*; *Aphis nerii* was estimated with *Aphis gossypii*; *Procambarus virginalis* was estimated with *Procambarus clarkii*; and *Pinctada fucata* was estimated with *Pinctada imbricata*. Otherwise, they were estimated with a species from the same family: *Echinus esculentus* was estimated with *Strongylocentrotus intermedius*; and *Parasteatoda tepidariorum* was estimated with *Theridion posticatum*. Ancestral state reconstruction was done in R using the function *make.simmap* from phytools 2.4.4^117^, using an equal rates (ER) model and 1000 simulations. We used the simple ER models, as all rates different (ARD) models had ι1AIC < 2 for both traits.

## Acknowledgements

We would like to thank Henri Truchassout and Laura Zamorano for discussion, and Romain Villoutreix for helping prepare samples for sequencing.

## Notes

https://github.com/planidin/epigen_sims

### Competing Interest Statement

The authors have declared no competing interest.

### Funding

- European Research Council: Grant agreement No.770826 EE-Dynamics
- Laboratoire d'Excellence TULIP: Senior Package to P.N.

