## Supplementary Information for "Dual roles of DNA methylation in gene expression and chromatin structure in a stick insect"

### Supplementary Results

*TSS versus gene body methylation*

Statistical tests comparing the 5% most and 5% least expressed genes are given in the main text (Figure S1). Here we compare the 25% most expressed and 25% least expressed genes (Figure S2). In highly expressed genes, the TSS had less methylation than unexpressed genes (6.8% vs 12.9% methylation, respectively; Wilcoxon rank-sum test: $n_1 = 1597$, $n_2 = 1553$, $W = 1490610$, $p < 1\times10^{-16}$; Figure S2A). The TSS was also more likely to contain a peak of open chromatin in highly expressed genes (85% of TSSs with an open chromatin peak vs 31% in unexpressed genes; Wilcoxon rank-sum test: $n_1 = 1597$, $n_2 = 1586$, $W = 580026$, $p < 1\times10^{-16}$; Figure S2B). In contrast, the gene body of highly expressed genes had substantially increased methylation (47.5% vs 21.8% in unexpressed genes; Wilcoxon rank-sum test: $n_1 = 1663$, $n_2 = 1238$, $W = 610817$, $p < 1\times10^{-16}$; Figure S2A), and fewer open chromatin peaks (0.26 vs 0.81 open chromatin peaks per kilobase in unexpressed genes; Wilcoxon rank-sum test: $n_1 = 1663$, $n_2 = 1283$, $W = 741486$, $p < 1\times10^{-16}$; Figure S2B).

The same trend is apparent when examining expression deciles (Figure S3). Among expression deciles, differences in TSS methylation (Kruskal-Wallis rank sum test: $n = 6338$, $\chi^2_{10} = 398$, $p < 1\times10^{-16}$; Figure S3A), gene body methylation (Kruskal-Wallis rank sum test: $n = 6180$, $\chi^2_{10} = 694$, $p < 1\times10^{-16}$; Figure S3A), open chromatin at the TSS (Pearson's $\chi^2$ test: $n = 6371$, $\chi^2_{10} = 1348$, $p < 1\times10^{-16}$; Figure S3B), and peaks per kilobase in the gene body (Kruskal-Wallis rank sum test: $n = 6230$, $\chi^2_{10} = 372$, $p < 1\times10^{-16}$; Figure S3B) all being statistically significant. However, the relationship between methylation and expression is not perfectly monotonic. For example, TSS methylation is lower for unexpressed genes than lowly expressed genes (1$^{st}$ decile), and gene body methylation is lower for highly expressed genes (10$^{th}$ decile) compared to moderately expressed genes (3$^{rd}$ to 8$^{th}$ deciles). This non-monotonicity can be in part explained by chromatin compartments. Unexpressed genes are more likely to occur in low methylation heterochromatin (Figure S3C), which would reduce methylation at their TSSs. Furthermore, highly expressed genes (10$^{th}$ decile) are less likely to occur in high methylation euchromatin than moderately expressed genes (3$^{rd}$ to 8$^{th}$ deciles; Figure S3C). This may be due to highly expressed genes occurring in regions of facultative rather than constitutive euchromatin.

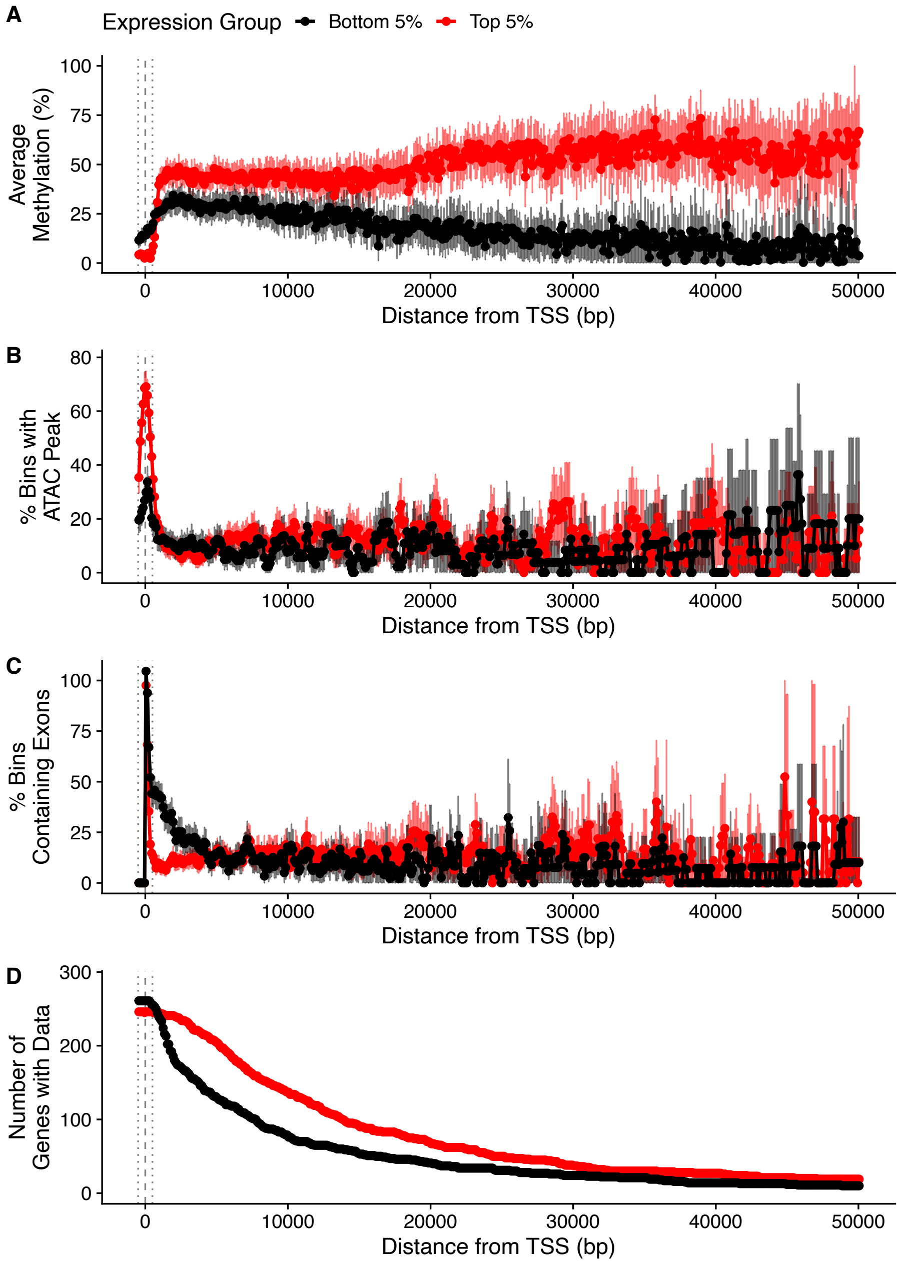


**Figure S1.** Comparison of the patterns of epigenetic marks along the gene bodies of the most and least expressed genes ($95^{th}$ versus $5^{th}$ quantiles). (A) DNA methylation, (B) ATAC-seq peak presence, (C) exon content, and (D) sample size for 100 bp bins based on distance from the transcription start site (TSS). Points are the average for each bin and error bars are a 95% confidence interval for the average. The vertical dashed line indicates the TSS, and the vertical dotted lines indicate 500 bp upstream and downstream of the TSS, representing the putative TSS-adjacent region.

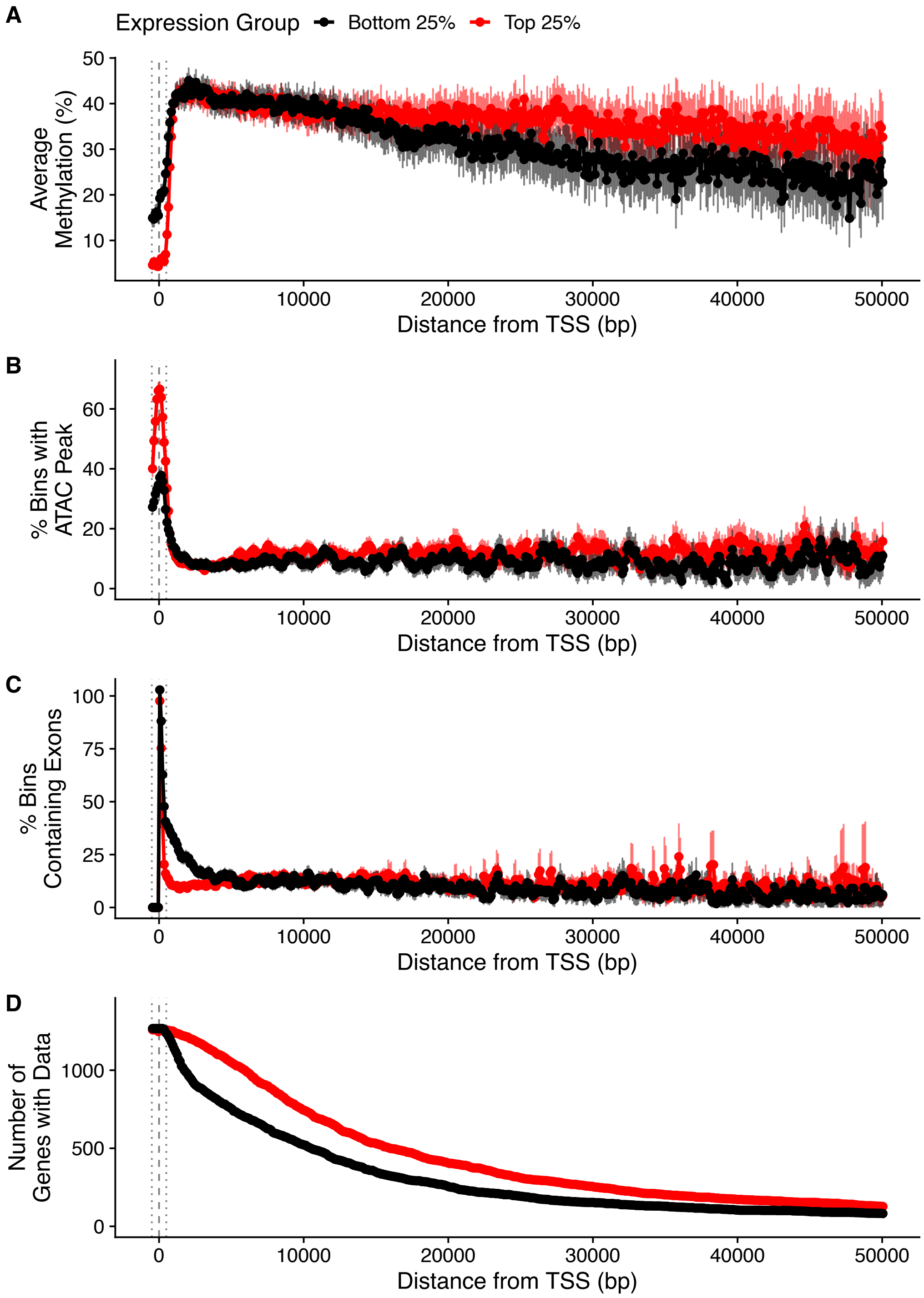


**Figure S2.** Comparison of the patterns of epigenetic marks along the gene bodies of the most and least expressed genes (75th versus 25th quantiles). (A) DNA methylation, (B) ATAC-seq peak presence, (C) exon content, and (D) sample size for 100 bp bins based on distance from the transcription start site (TSS). Points are the average for each bin and error bars are a 95% confidence interval for the average. The vertical dashed line indicates the TSS, and the vertical dotted lines indicate 500 bp upstream and downstream of the TSS, representing the putative TSS-adjacent region.

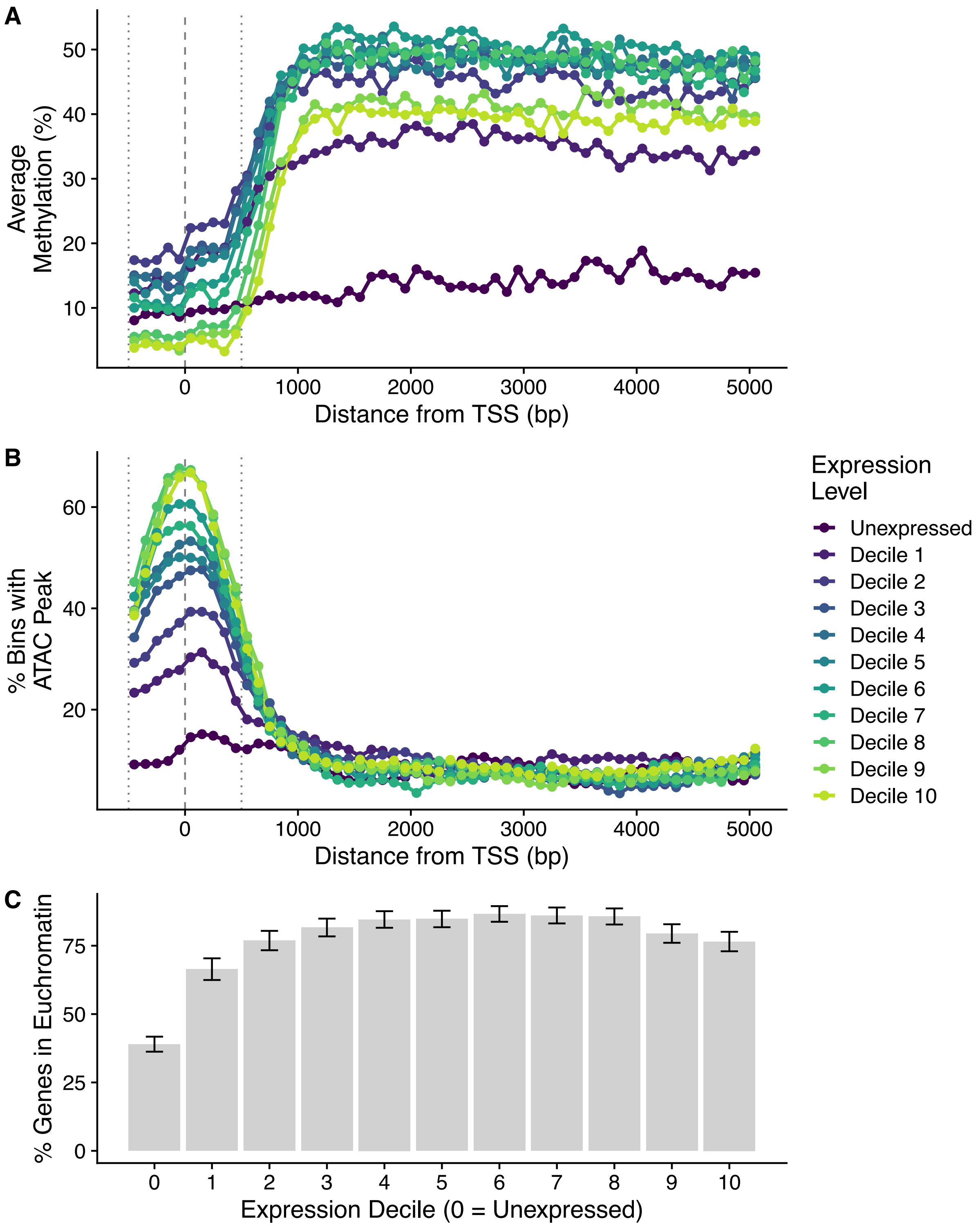


**Figure S3.** Comparison of the patterns of epigenetic marks along the gene bodies for genes binned by expression decile. (A) DNA methylation, (B) ATAC-seq peak presence. Points are the average for each bin. The vertical dashed line indicates the TSS, and the vertical dotted lines indicate 500 bp upstream and downstream of the TSS, representing the putative TSS-adjacent region. (C) Percent of genes in euchromatin by expression decile. Error bars show the 95% CI.

*Changes in methylation along the gene body*

The amount of gene body methylation varies along the gene body in invertebrates with the pattern depending on the species, and typically with greater methylation on exons [1–6]. However, our observations suggest that much of the variation in gene body methylation in *T. cristinae* is due to proximity to the TSS, rather than exon/intron number or gene length. Specifically, in *T. cristinae* gene body methylation was previously shown to be greater for later introns and exons until about the fifth exon where it began to decline [1]. However, here we find that differences in methylation between exons one through five are entirely due to the region of reduced methylation adjacent to the TSS, and the decrease in later exons is due to differences in methylation among, rather than within, genes. There is an inverse U-shaped relationship between gene length and methylation, whereas expression monotonically increases with gene length (Figure S4A-B). This makes bins further from the TSS appear to have less methylation, as shorter genes lack these bins (Figure S4C). In contrast, methylation residual to the average gene body methylation is roughly constant after the first ~1000 bp downstream of the TSS (Figure S4D). There is, however, an increase in methylation by about 5% right before the TES after normalization, likely due to increased exon content at the end of genes (Figure S4E-F). Lastly, exons remain elevated in methylation relative to introns, even when controlling for their position relative to the TSS (Figure S4G-H). These results suggest that the pattern of gene body methylation in *T. cristinae* are driven by changes at the TSS and variation among genes. This may also hold true for other invertebrates that have different relationships between gene length, expression and gene body methylation [7–9].

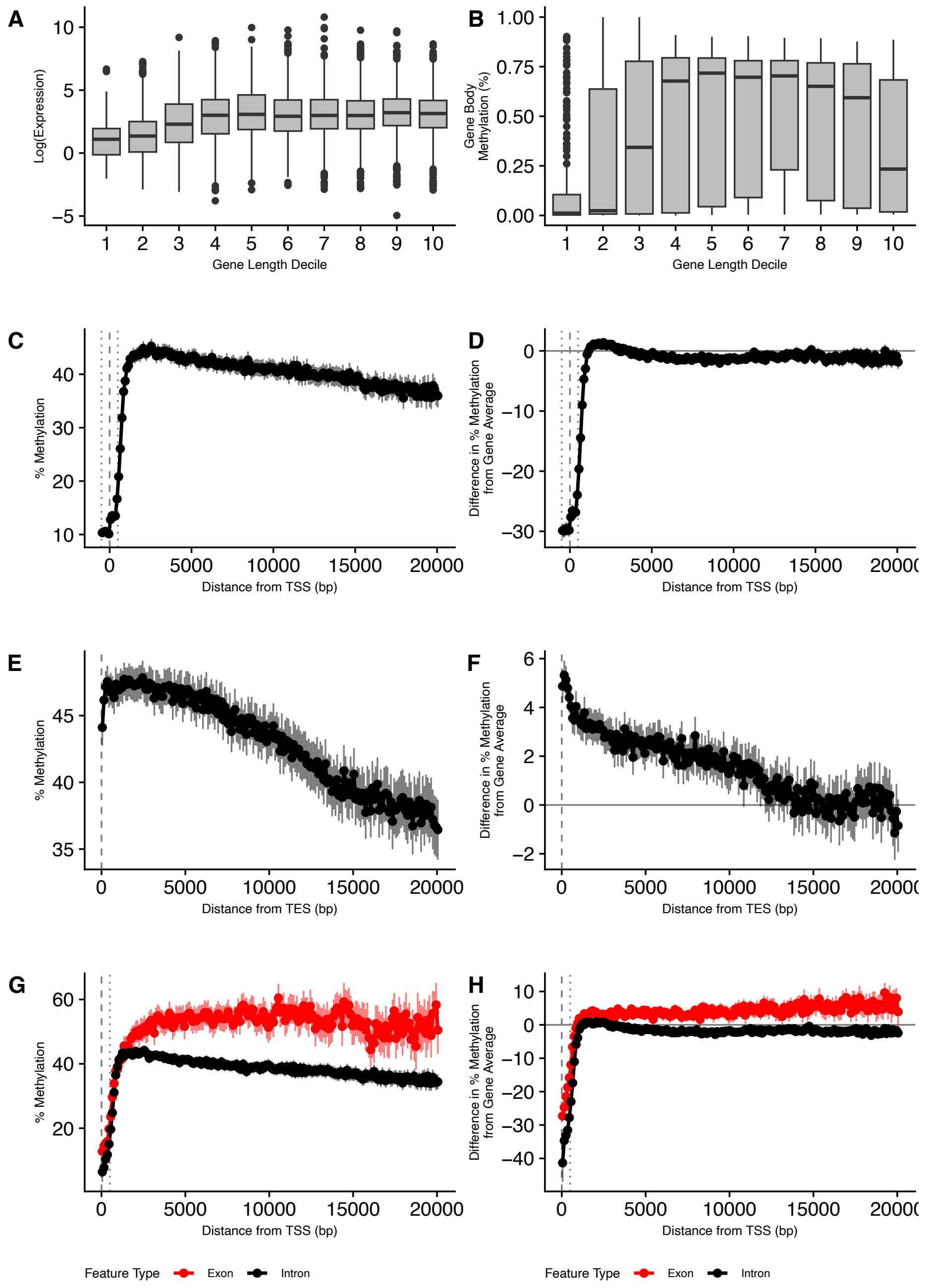


**Figure S4.** (A) Gene expression, and (B) methylation, for genes categorized by length deciles. (C) Gene body methylation in 100 bp bins compared to (D) gene body methylation residual to average gene body methylation, when aligned by the TSS. Average gene body methylation was calculated using bins more than 500 bp downstream of the transcription start site (TSS) (E) Gene body methylation in 100 bp bins compared to (F) gene body methylation residual to average gene body methylation, when aligned by the TES. Only bins at least 1000 bp downstream of the TSS were used to generate plots E and F. (G) Gene body methylation in exons and introns, compared to (H) gene body methylation residual to average gene body methylation. Points are the average for each bin and error bars are a 95% confidence interval for the average. The vertical dashed line indicates the TSS, and the vertical dotted lines indicate 500 bp upstream and downstream of the TSS, representing the putative TSS-adjacent region.

*Variation in methylation and expression between chromatin compartments*

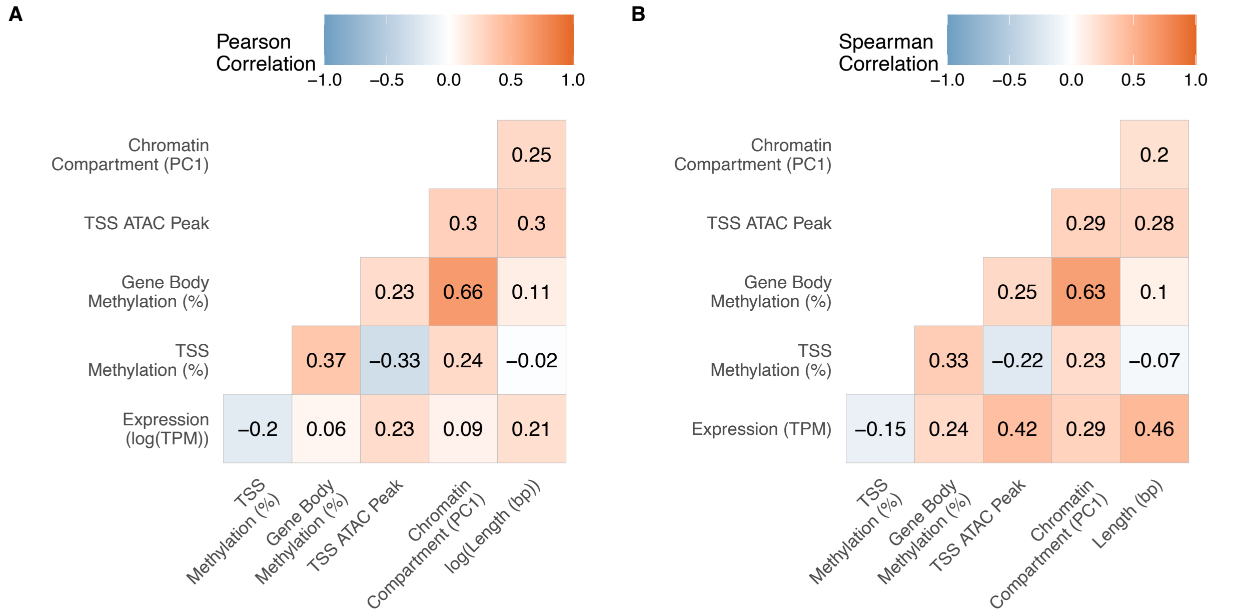


**Figure S5.** (A) Pearson correlation among epigenetic marks, gene expression and gene length. (B) Spearman correlation.

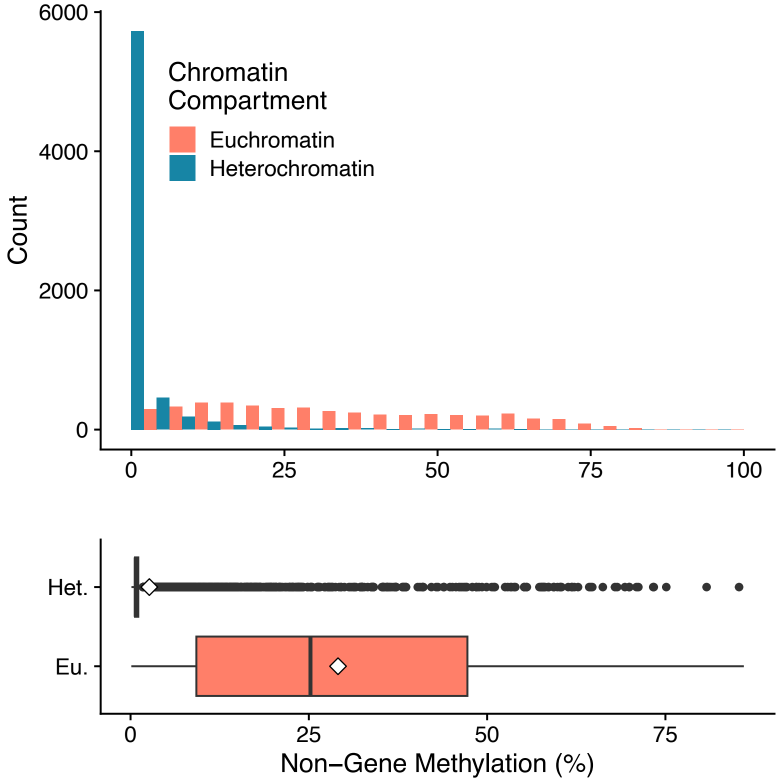


**Figure S6.** Histogram of average percent DNA methylation in 100kb bins for CpG sites outside of gene annotations, depending on whether the bin is in a region of putative euchromatin or heterochromatin. Boxplot shows the same data, with diamonds displaying the mean for each group.

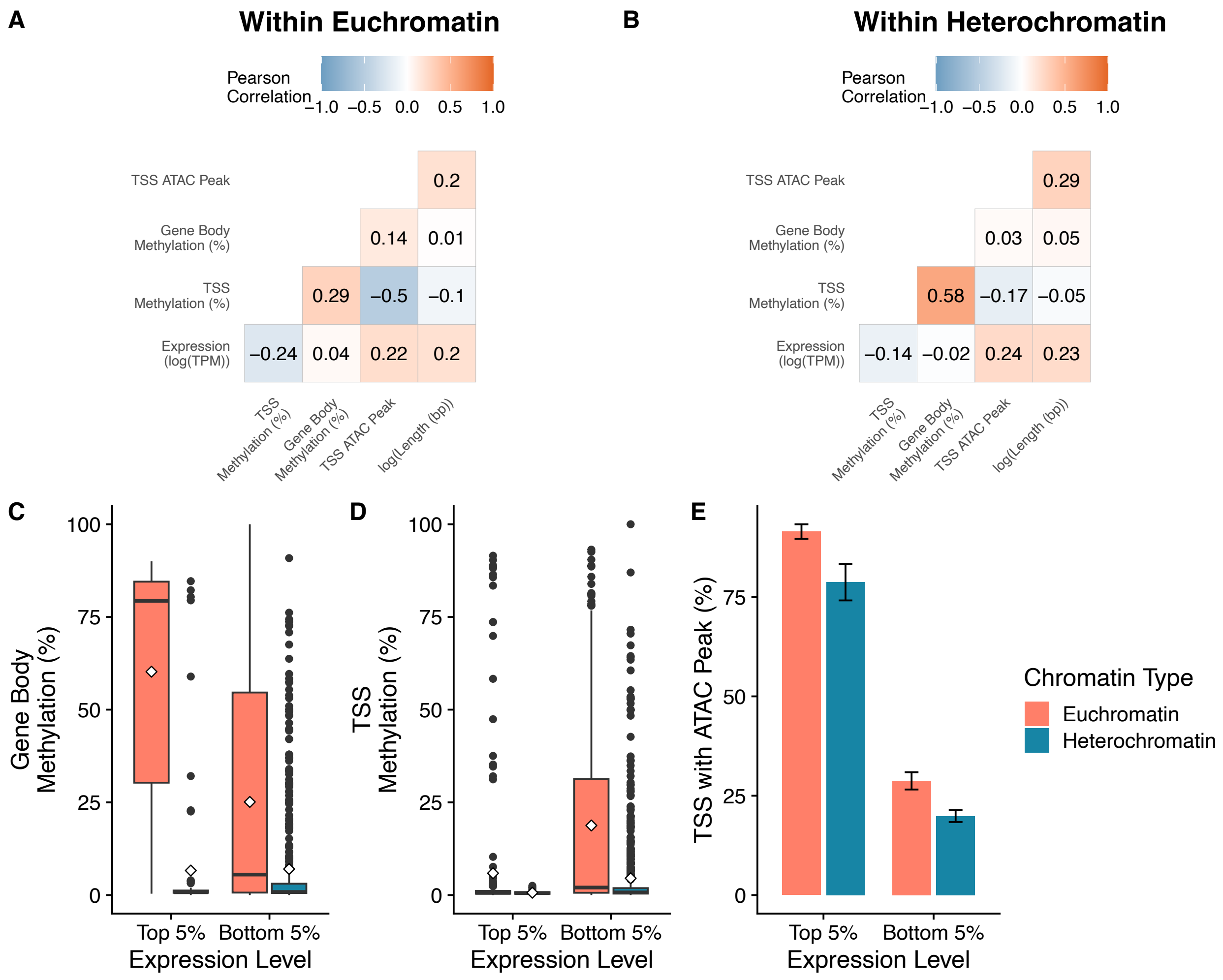


**Figure S7.** Pearson correlations for epigenetic marks, gene expression and gene length, for genes within (A) euchromatin or (B) heterochromatin compartments. Comparison of (B) transcription start site (TSS) methylation, (C) TSS openness, and (D) gene body methylation, for highly expressed and unexpressed genes in euchromatin and heterochromatin compartments.

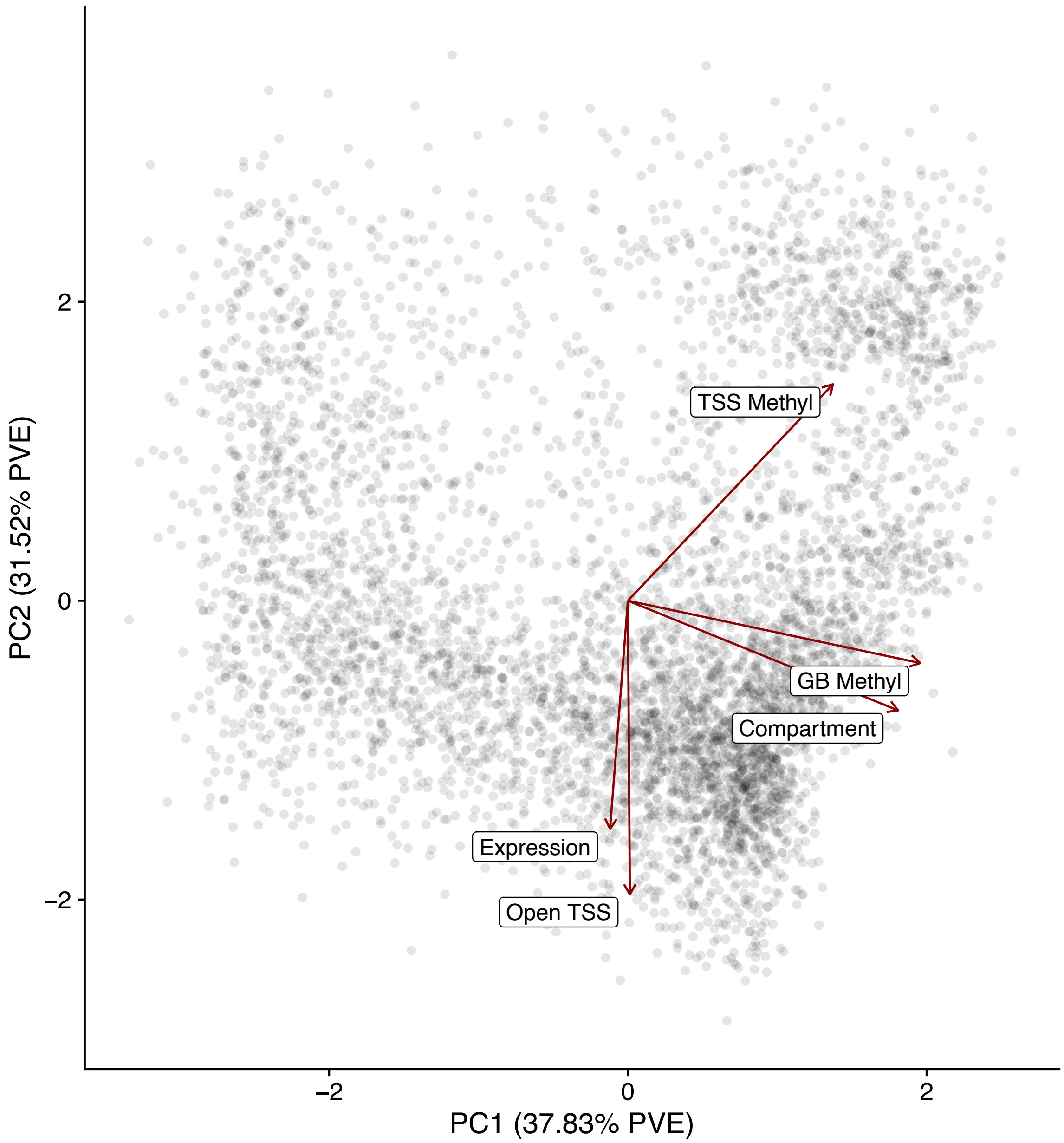


**Figure S8.** First two principal components of the 'epigenetic state' of genes, based on log(expression), logit(gene body (GB) methylation), logit(transcription start site (TSS) methylation), chromatin compartment score (PC1), and presence of an open chromatin peak at the TSS. Arrows show variable loadings, and each point is an individual gene.

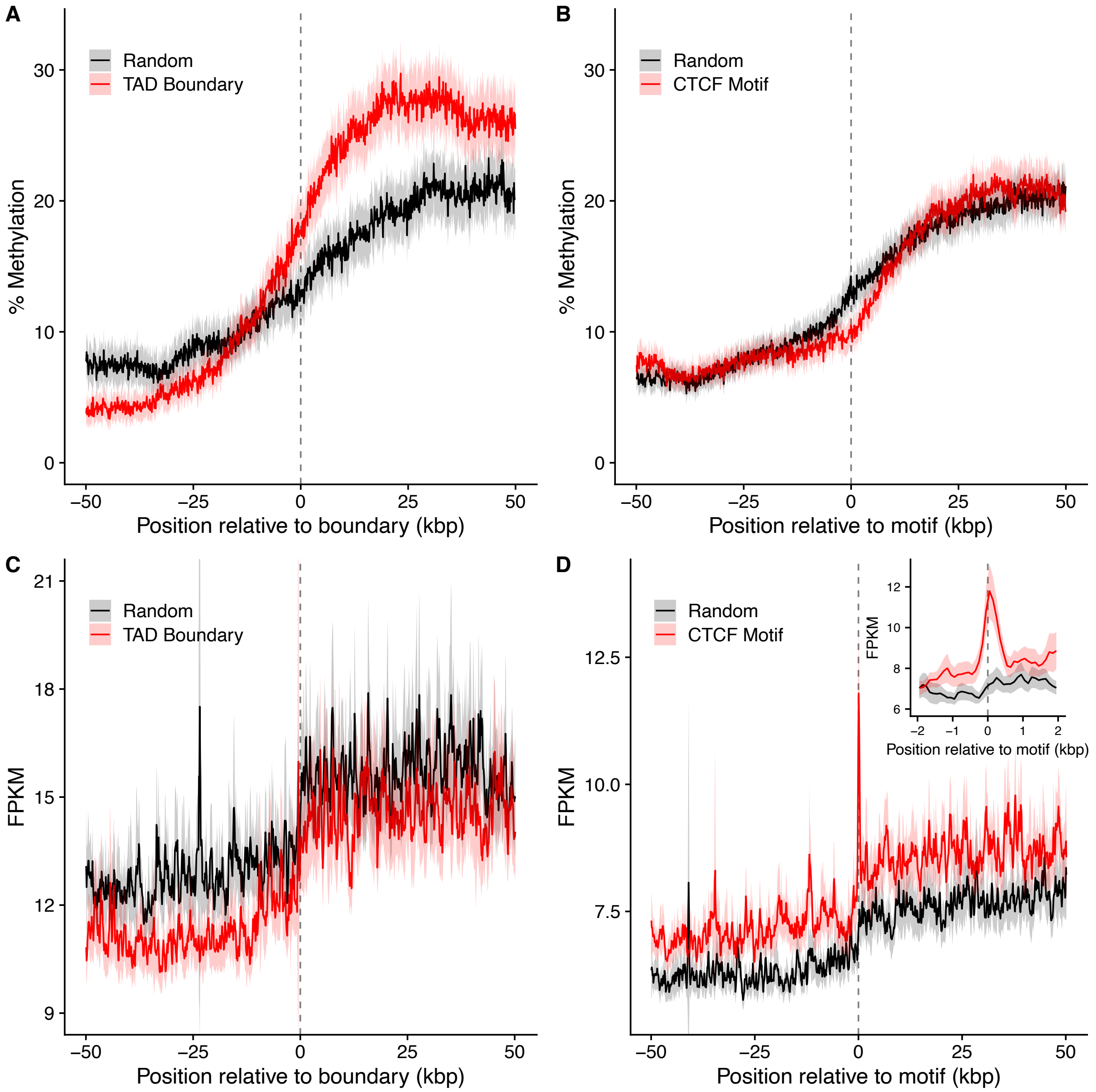


**Figure S9.** Profiles of (A, B) % methylation and (C, D) ATAC-seq fragments per kilobase per million fragments (FPKM) at (A, C) topologically associated domain (TAD) boundaries and (B, D) 'CCCTC-binding factor (CTCF)'-binding motifs. Both % methylation and FPKM were summarized in 100 bp bins. Lines show the average among all TAD boundaries or CTCF-binding motifs. Filled regions give the 95% CI for the mean. Random data was generated by randomly selecting the same number of points in the genome as there was a given motif. Data at each site was oriented to have the side with the greater average on the right, prior to averaging among sites for both real and randomized data.

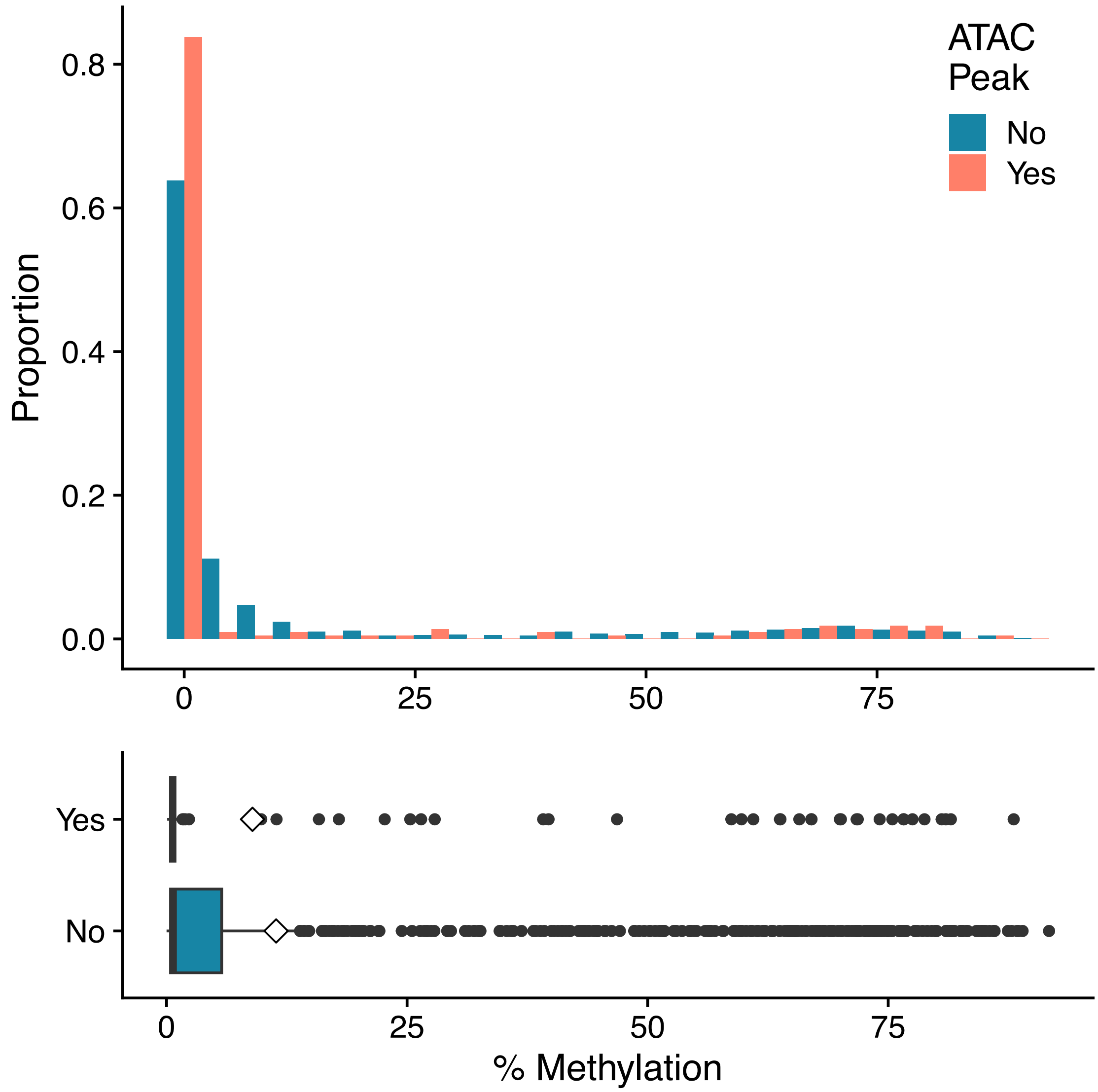


**Figure S10.** Histogram of average percent DNA methylation for the 500 bp surrounding CTCF-binding motifs, depending on whether the motif overlaps an ATAC-seq peak. Boxplot shows the same data, with diamonds displaying the mean for each group.

*Comparative analysis of* Apis mellifera

Statistical tests comparing the 5% most and 5% least expressed genes are given in the main text (Figure S12-13). Here we compare the 25% most expressed and 25% least expressed genes (Figure S14). The TSS had slightly more methylation in highly expressed genes than in unexpressed genes (1.8% vs 1.7% methylation, respectively; Wilcoxon rank-sum test: $n_1 = 1249$, $n_2 = 1447$, $W = 831364.5$, $p = 3\times10^{-4}$; Figure S14A). The TSS was also more likely to contain a peak of open chromatin in highly expressed genes (43% of TSSs with an open chromatin peak vs 29% in unexpressed genes; Wilcoxon rank-sum test: $n_1 = 1252$, $n_2 = 1464$, $W = 782428$, $p = 2\times10^{-15}$; Figure S14B). In contrast, the gene body of highly expressed genes had substantially increased methylation (30.0% vs 4.4% in unexpressed genes; Wilcoxon rank-sum test: $n_1 = 1692$, $n_2 = 1541$, $W = 468458$, $p < 1\times10^{-16}$; Figure S14A), and marginally fewer open chromatin peaks per kilobase compared to low expression genes (0.17 vs 0.21 open chromatin peaks per kilobase, respectively; Wilcoxon rank-sum test: $n_1 = 1713$, $n_2 = 1556$, $W = 1379290$, $p = 0.052$; Figure S14B).

The same trend is apparent when examining expression deciles (Figure S15). Among expression deciles, differences in TSS methylation (Kruskal-Wallis rank sum test: $n = 5147$, $\chi^2_{10} = 122$, $p < 1\times10^{-16}$; Figure S145), gene body methylation (Kruskal-Wallis rank sum test: $n = 6641$, $\chi^2_{10} = 1542$, $p < 1\times10^{-16}$; Figure S15A), open chromatin at the TSS (Pearson's $\chi^2$ test: $n = 5169$, $\chi^2_{10} = 122$, $p < 1\times10^{-16}$; Figure S15B), and peaks per kilobase in the gene body (Kruskal-Wallis rank sum test: $n = 6707$, $\chi^2_{10} = 37$, $p = 7\times10^{-5}$; Figure S15B) all being statistically significant. The relationship between methylation and expression in *A. mellifera* is more monotonic than in *T. cristinae*, but gene body methylation is lower for highly expressed genes (10$^{th}$ decile) compared to moderately expressed genes (4$^{th}$ to 8$^{th}$ deciles). However, highly expressed genes (10$^{th}$ decile) are not less likely to occur in high methylation euchromatin than moderately expressed genes (4$^{th}$ to 8$^{th}$ deciles; Figure S15C).

Our observations suggest that differences in gene body methylation with gene length in *A. mellifera* is due to a peak of methylation around 1000 bp from the TSS. There is a negative relationship between gene length and methylation, except for the least expressed genes, whereas there is no relationship between gene length and gene expression (Figure S16A-B). This is caused by a peak of methylation around 1000 bp from the TSS that makes up a smaller portion of longer genes, causing longer genes to have overall lower methylation (Figure S16C). This pattern remains when looking at methylation residual to the average gene body methylation (Figure S16D). While there is an increase in methylation towards the TES (Figure S16E), there is actually a decline in methylation towards the TES when measuring methylation residual to the gene body average (Figure S16F), likely due to the same peak 1000 bp from the TSS. Lastly, exons are elevated in methylation relative to introns within the 'peak' region, but not along the rest of the gene body, when measuring methylation residual to the gene body average (Figure S16G-H). These results suggest that variation in gene body methylation in *A. mellifera* is driven by a small region around 1000 bp downstream of the TSS.

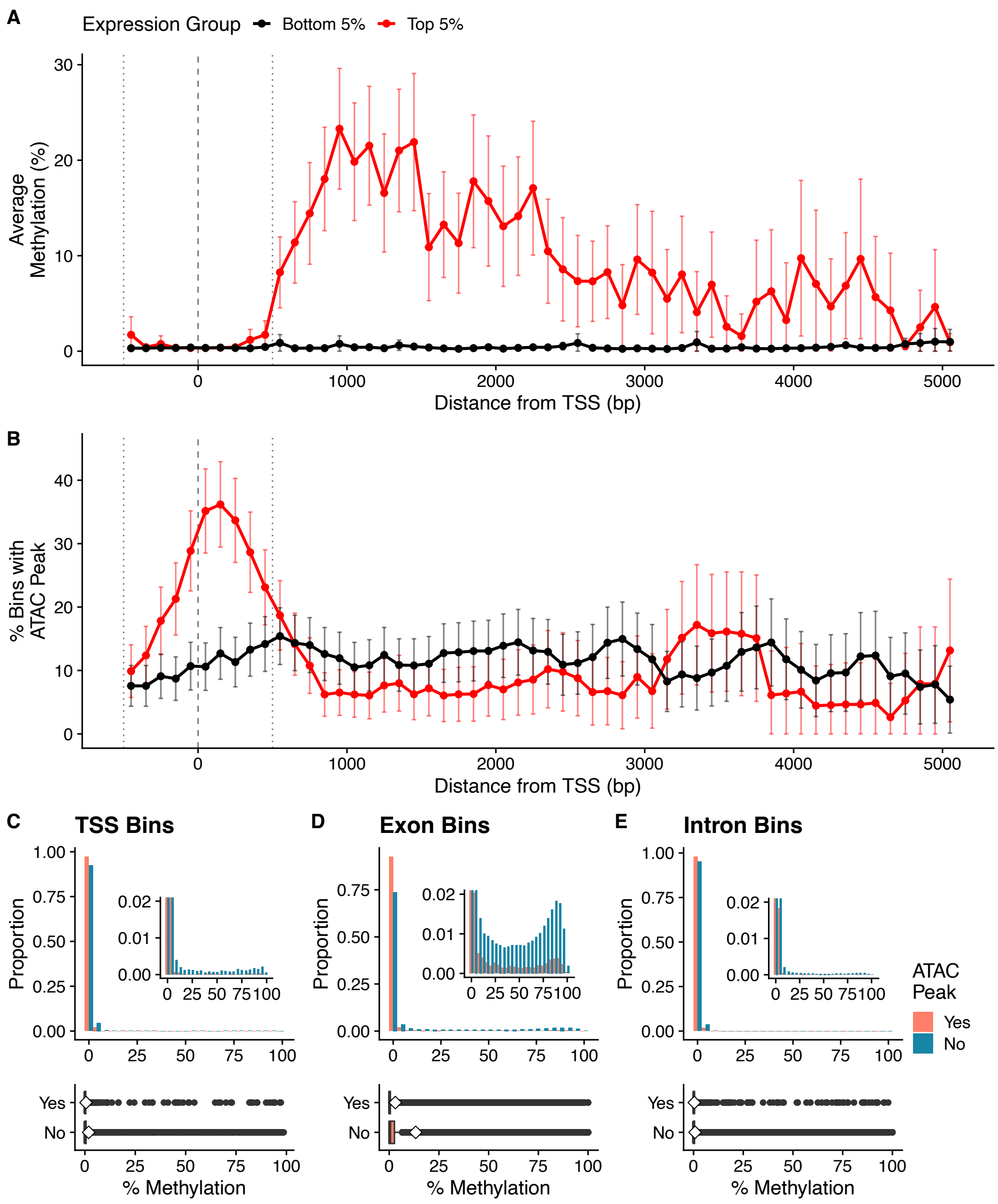


**Figure S11.** (A) In *Apis mellifera*, there is no difference in methylation at the transcription start site (TSS) between highly and unexpressed genes, but there is an increased gene body methylation, along with (B) a greater probability of having an open chromatin peak at their TSS. This relationship is modulated by the repression of open chromatin peaks by methylation at (C) the TSS, (D) exons, and (E) introns. (A-B) Points are averages for a 100 bp bin, across all genes within a given expression class, and error bars are the 95% CI. The vertical dashed line indicates the TSS, and the vertical dotted lines indicate the 'TSS-adjacent' region used in subsequent analyses. (C-E) Histograms show the percent methylation for 100 bp bins along all genes, dividing between bins containing and not containing an ATAC-seq peak. Boxplots show the same data, displaying median (center line), quartiles (box), and quartiles plus 1.5 times the interquartile range (whiskers), with diamonds indicating the average. (C) Exonic and intronic bins within 500 bp of the TSS. (D) Exonic bins more than 500 bp from the TSS. (E) Intronic bins more than 500 bp from the TSS.

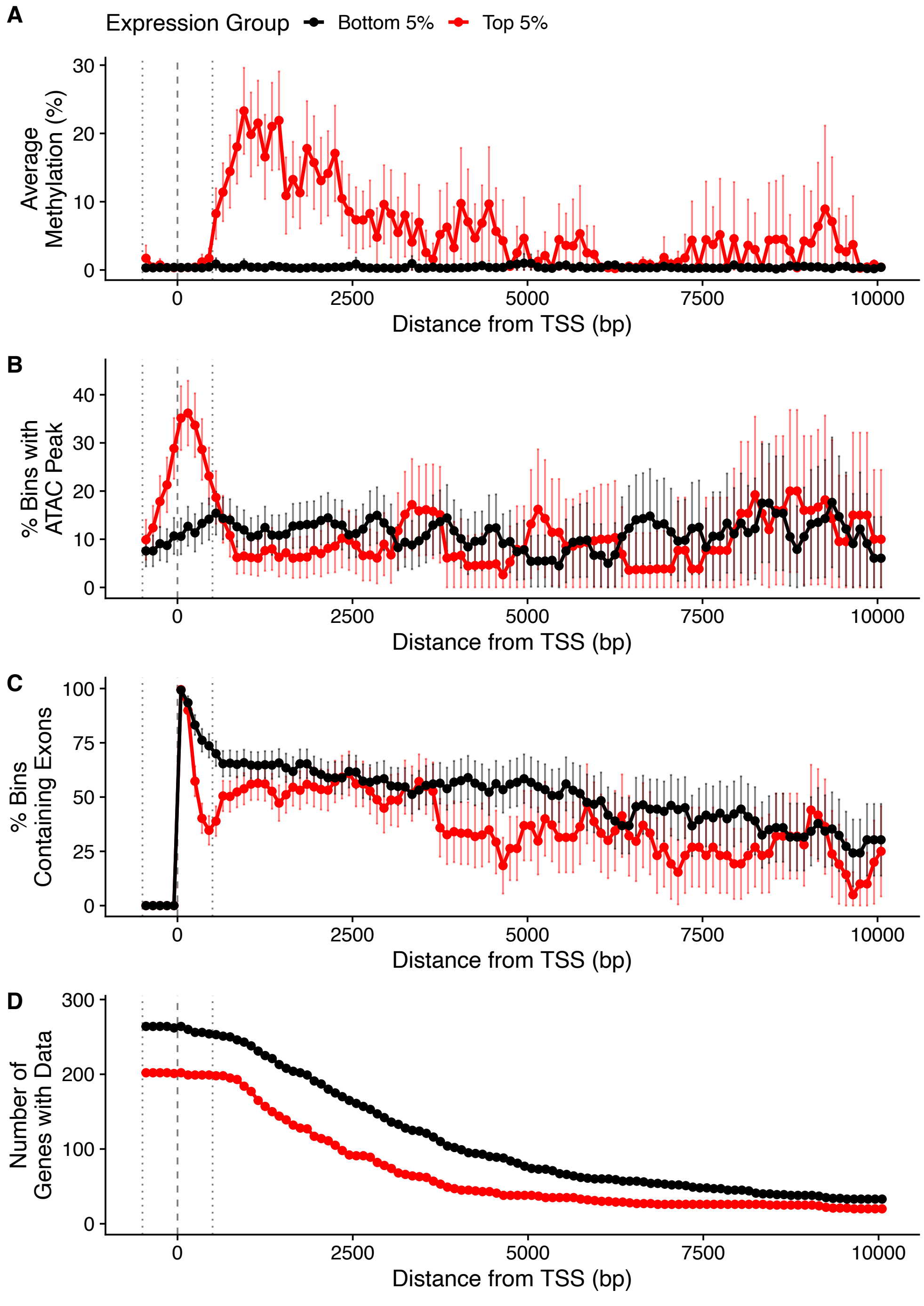


**Figure S12.** Comparison of the patterns of epigenetic marks along the gene bodies of the most and least expressed genes (95th versus 5th quantiles) in *Apis mellifera*. (A) DNA methylation, (B) ATAC-seq peak presence, (C) exon content, and (D) sample size for 100 bp bins based on distance from the transcription start site (TSS). Points are the average for each bin and error bars are a 95% confidence interval for the average. The vertical dashed line indicates the TSS, and the vertical dotted lines indicate 500 bp upstream and downstream of the TSS, representing the putative TSS-adjacent region.

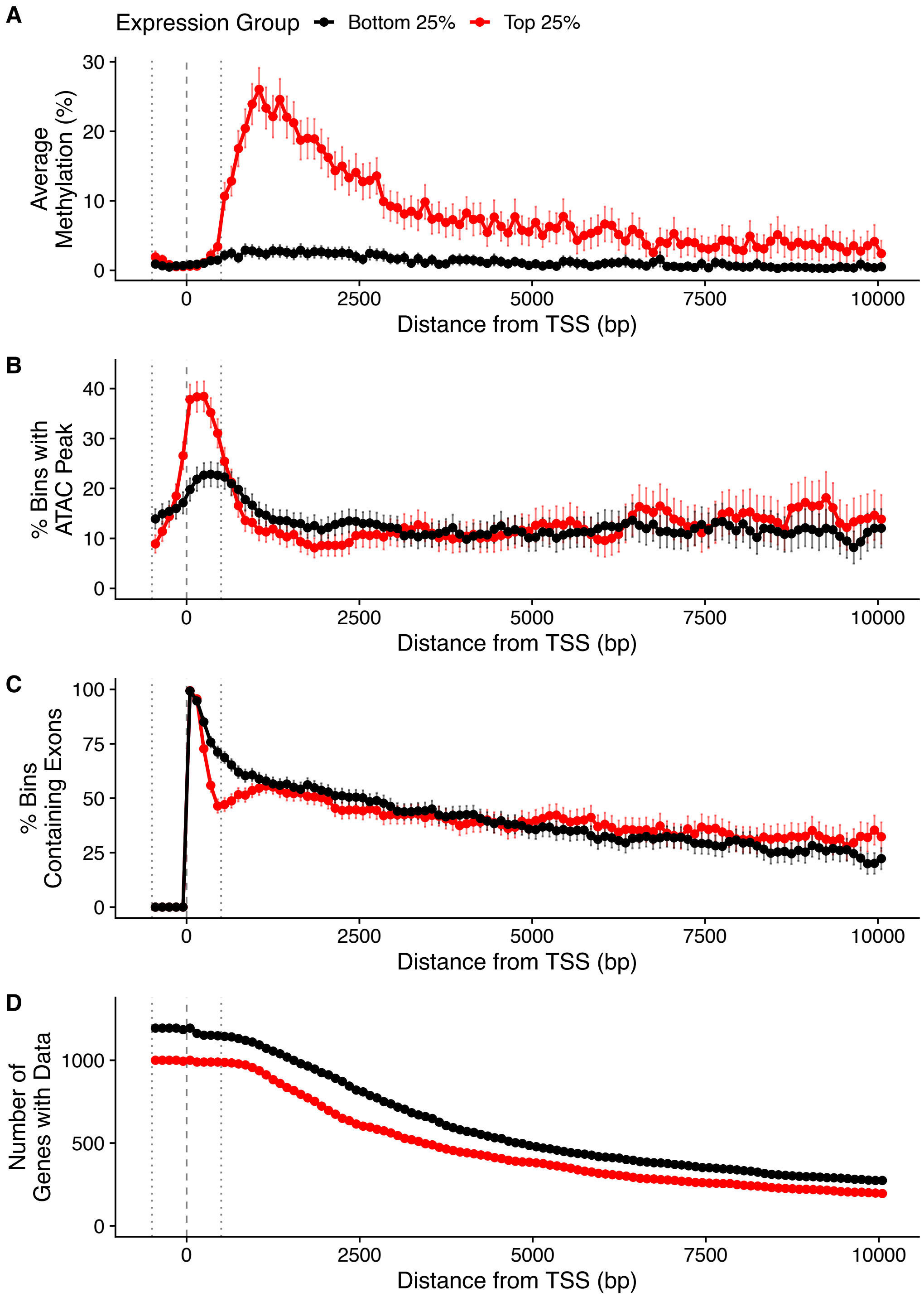


**Figure S13.** Comparison of the patterns of epigenetic marks along the gene bodies of the most and least expressed genes (75th versus 25th quantiles) in *Apis mellifera*. (A) DNA methylation, (B) ATAC-seq peak presence, (C) exon content, and (D) sample size for 100 bp bins based on distance from the transcription start site (TSS). Points are the average for each bin and error bars are a 95% confidence interval for the average. The vertical dashed line indicates the TSS, and the vertical dotted lines indicate 500 bp upstream and downstream of the TSS, representing the putative TSS-adjacent region.

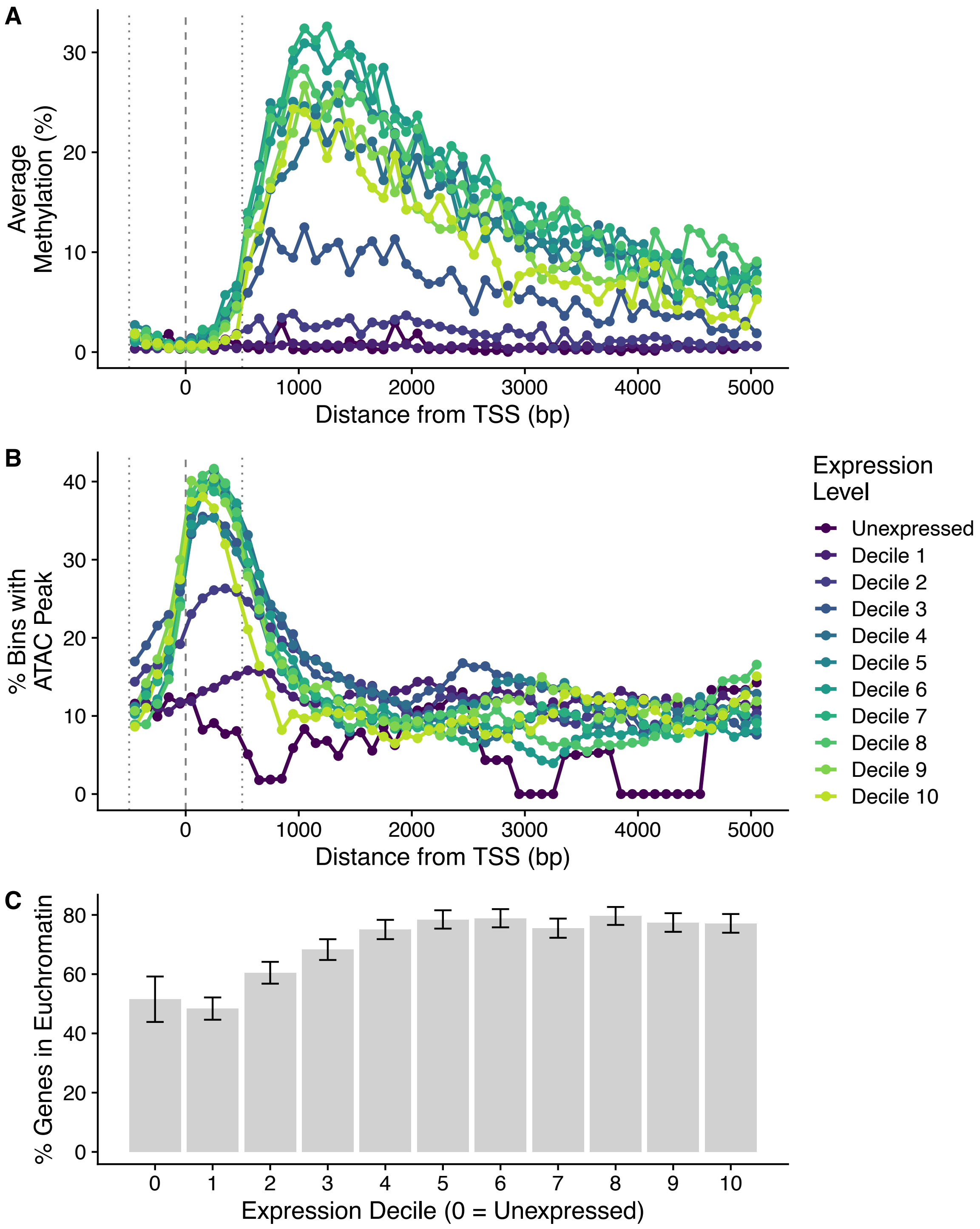


**Figure S14.** Comparison of the patterns of epigenetic marks along the gene bodies for genes binned by expression decile in *Apis mellifera*. (A) DNA methylation, (B) ATAC-seq peak presence. Points are the average for each bin. The vertical dashed line indicates the TSS, and the vertical dotted lines indicate 500 bp upstream and downstream of the TSS, representing the putative TSS-adjacent region. (C) Percent of genes in euchromatin by expression decile. Error bars show the 95% CI.

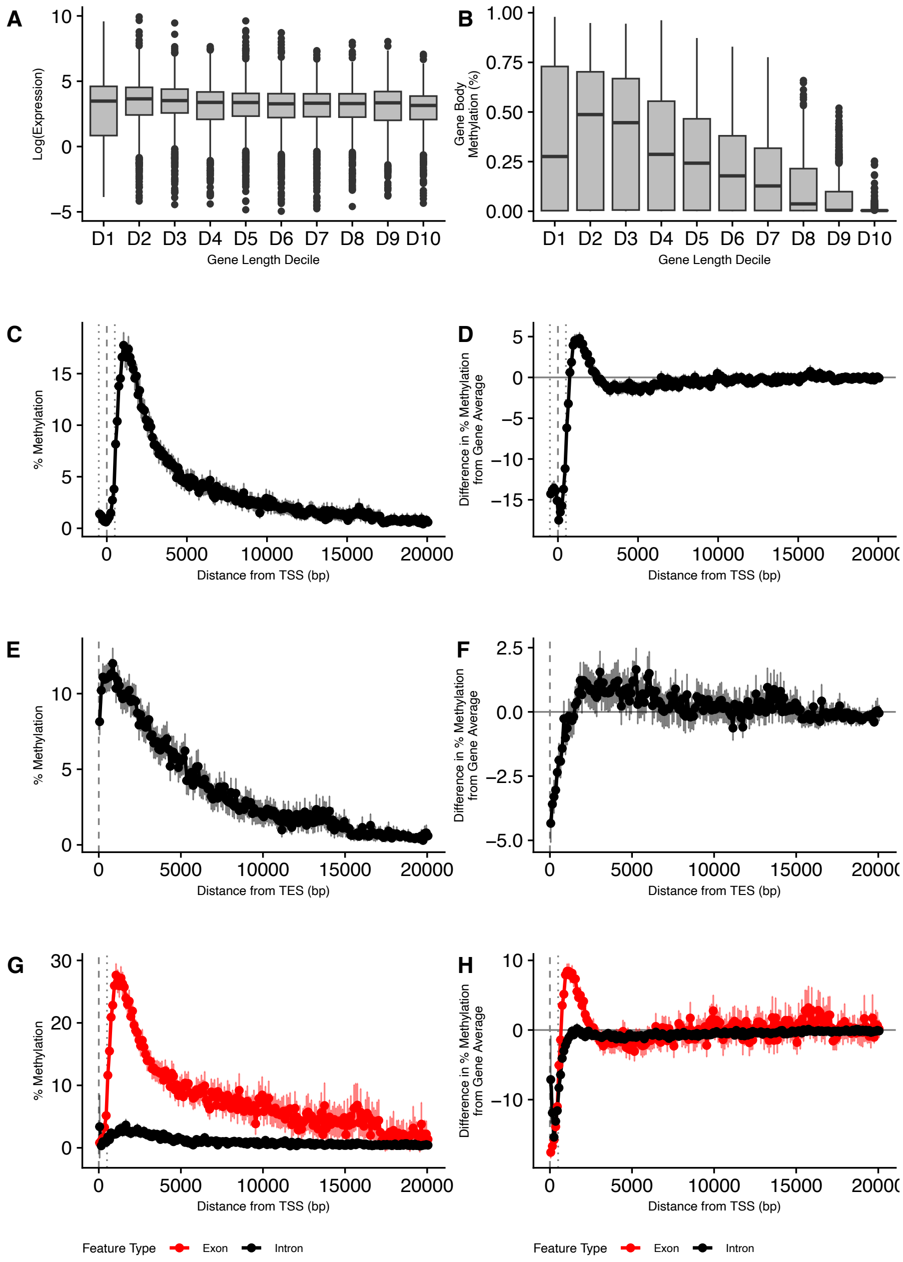


**Figure S15.** DNA methylation but not gene expression depends on gene length in *Apis mellifera* (A) Gene expression, and (B) methylation, for genes categorized by length deciles. (C) Gene body methylation in 100 bp bins compared to (D) gene body methylation residual to average gene body methylation, when aligned by the TSS. Average gene body methylation was calculated using bins more than 500 bp downstream of the transcription start site (TSS) (E) Gene body methylation in 100 bp bins compared to (F) gene body methylation residual to average gene body methylation, when aligned by the TES. Only bins at least 1000 bp downstream of the TSS were used to generate plots E and F. (G) Gene body methylation in exons and introns, compared to (H) gene body methylation residual to average gene body methylation. Points are the average for each bin and error bars are a 95% confidence interval for the average. The vertical dashed line indicates the TSS, and the vertical dotted lines indicate 500 bp upstream and downstream of the TSS, representing the putative TSS-adjacent region.

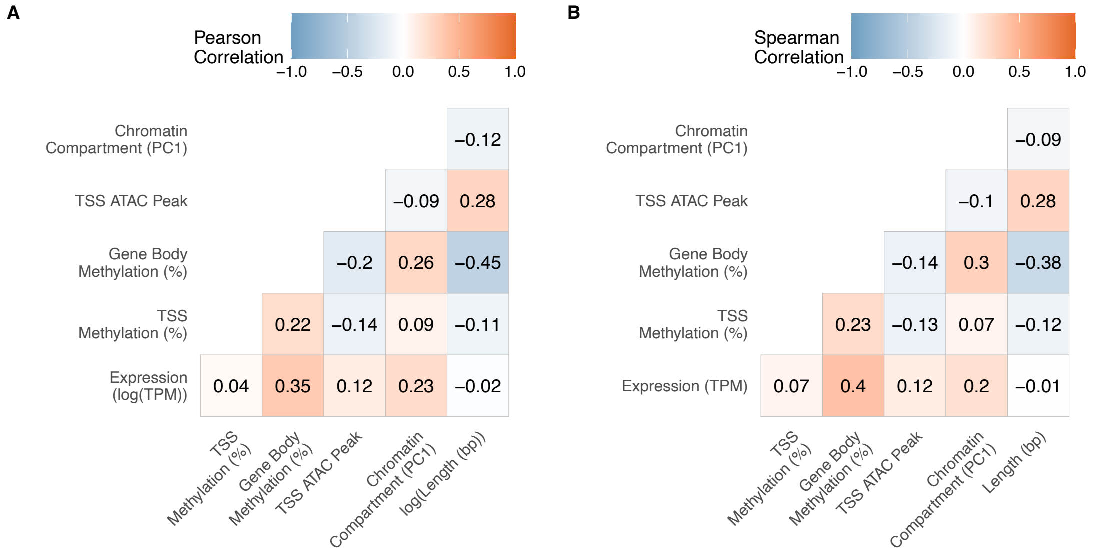


**Figure S16.** (A) Pearson correlation among epigenetic marks, gene expression and gene length in *Apis mellifera*. (B) Spearman correlation.

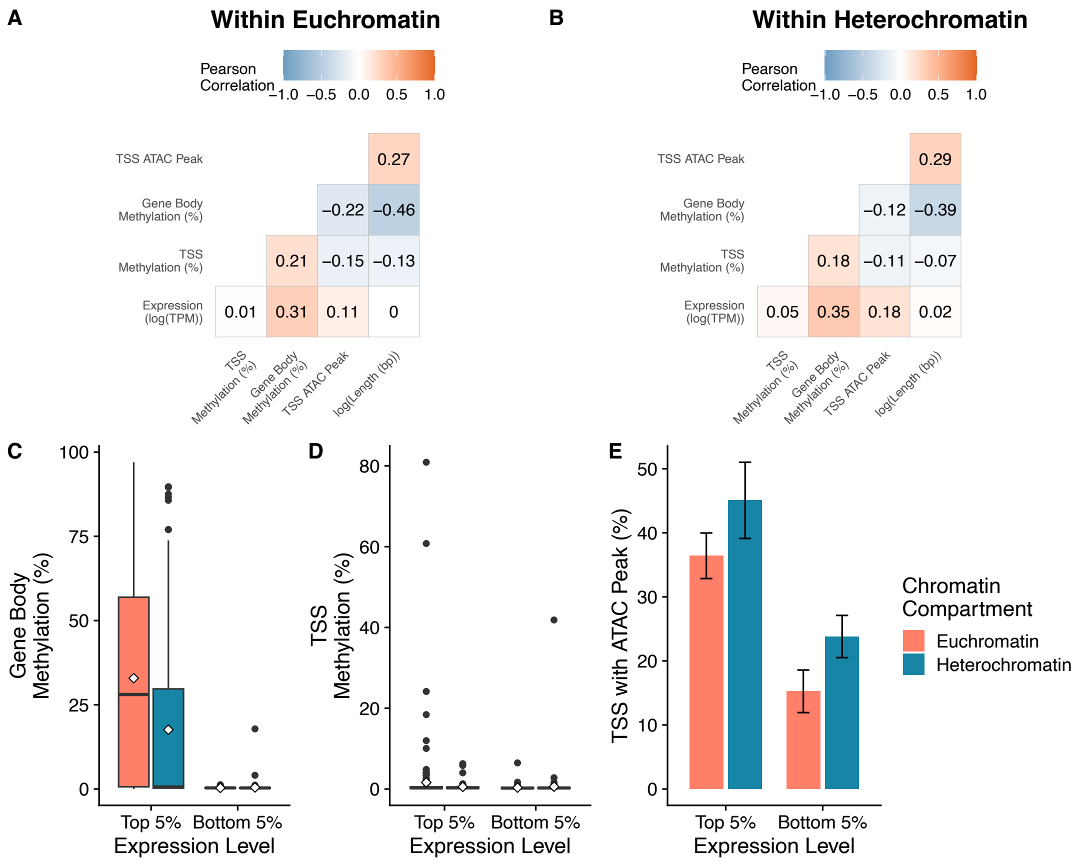


**Figure S17.** Pearson correlations for epigenetic marks, gene expression and gene length, for genes within (A) euchromatin or (B) heterochromatin compartments in *Apis mellifera*. Comparison of (B) transcription start site (TSS) methylation, (C) TSS openness, and (D) gene body methylation, for highly expressed and unexpressed genes in euchromatin and heterochromatin compartments.

**Table S1.** Species and references used to generate Figure 7. Transcription start site methylation was defined as a negative association between gene expression and DNA methylation at the promoter or regions flanking the TSS. Euchromatin methylation was defined as elevated methylation in euchromatin compartments relative to heterochromatin compartments. Species completely lacking DNA methylation were included for TSS methylation but not euchromatin methylation. Stars indicate species where euchromatin methylation is inferred from a positive association between DNA methylation and histone modifications associated with active chromatin.

| Species | TSS Methylation | Reference | Euchromatin Methylation | Reference |
|---|---|---|---|---|
| *Acyrthosiphon pisum* | 0 | Lewis et al.[10] | | |
| *Aphis nerii* | | | 1 | Mandrioli et al.[27] |
| *Apis mellifera* | 0 | This study; Lyko et al.[11] | 1 | This study; Hunt et al.[28] |
| *Arabidopsis thaliana* | 1 | Zhang et al.[12] | 0 | Wang et al.[29] |
| *Bombus terrestris* | 0 | Lewis et al.[10] | | |
| *Bombyx mori* | 0 | Xiang et al.[13] | 1* | Nanty et al.[30] |
| *Caenorhabditis elegans* | 0 | Simpson et al.[14] | | |
| *Camponotus floridanus* | | | 1* | Glastad et al.[31] |
| *Ciona intestinalis* | 1 | Keller et al.[3] | 1* | Nanty et al.[30] |
| *Crassostrea gigas* | 1 | Riviere et al.[15] | | |
| *Danio rerio* | 1 | Gibbs et al.[16] | 0 | Lindeman et al.[32] |
| *Drosophila melanogaster* | 0 | Rae & Steele[17] | | |
| *Echinus esculentus* | | | 1 | Bird et al.[33] |
| *Gallus gallus* | 1 | McGhee and Ginder[18] | 0 | Razin and Cedar[34] |
| *Heliconius melpomene* | 0 | Lewis et al.[10] | | |
| *Homo sapiens* | 1 | Busslinger et al.[19] | 0 | Lubit et al.[35] |
| *Limulus polyphemus* | 0 | Lewis et al.[10] | | |
| *Mus musculus* | 1 | Jahner et al.[20] | 0 | Miller et al.[36] |
| *Nasonia vitripennis* | 0 | Wang et al.[8] | | |
| *Neurospora crassa* | 0 | Bewick et al.[21] | 0 | Rountree and Selker[37] |
| *Nicrophorus vespilloides* | 0 | Lewis et al.[10] | | |
| *Parasteatoda tepidariorum* | 0 | Lewis et al.[10] | | |
| *Pinctada fucata martensii* | 1 | Zhang et al.[22] | | |
| *Planococcus citri* | 1 | Lewis et al.[10] | 1 | Bongiorni et al.[38] |
| *Procambarus virginalis* | 1 | Gatzmann et al.[23] | | |
| *Solenopsis invicta* | | | 1* | Hunt et al.[28] |
| *Strigamia maritima* | 1 | Lewis et al.[10] | | |
| *Strongylocentrotus* | 1 | Bogan et al.[24] | | |
| *Tetraodon nigroviridis* | 1 | Anastasiadi et al.[25] | | |
| *Timema cristinae* | 1 | This study | 1 | This study |
| *Xenopus tropicalis* | 1 | Ben-Hattar and Jiricny[26] | 0 | Bennet et al.[39] |
| *Zootermopsis nevadensis* | | | 1* | Glastad et al.[2] |

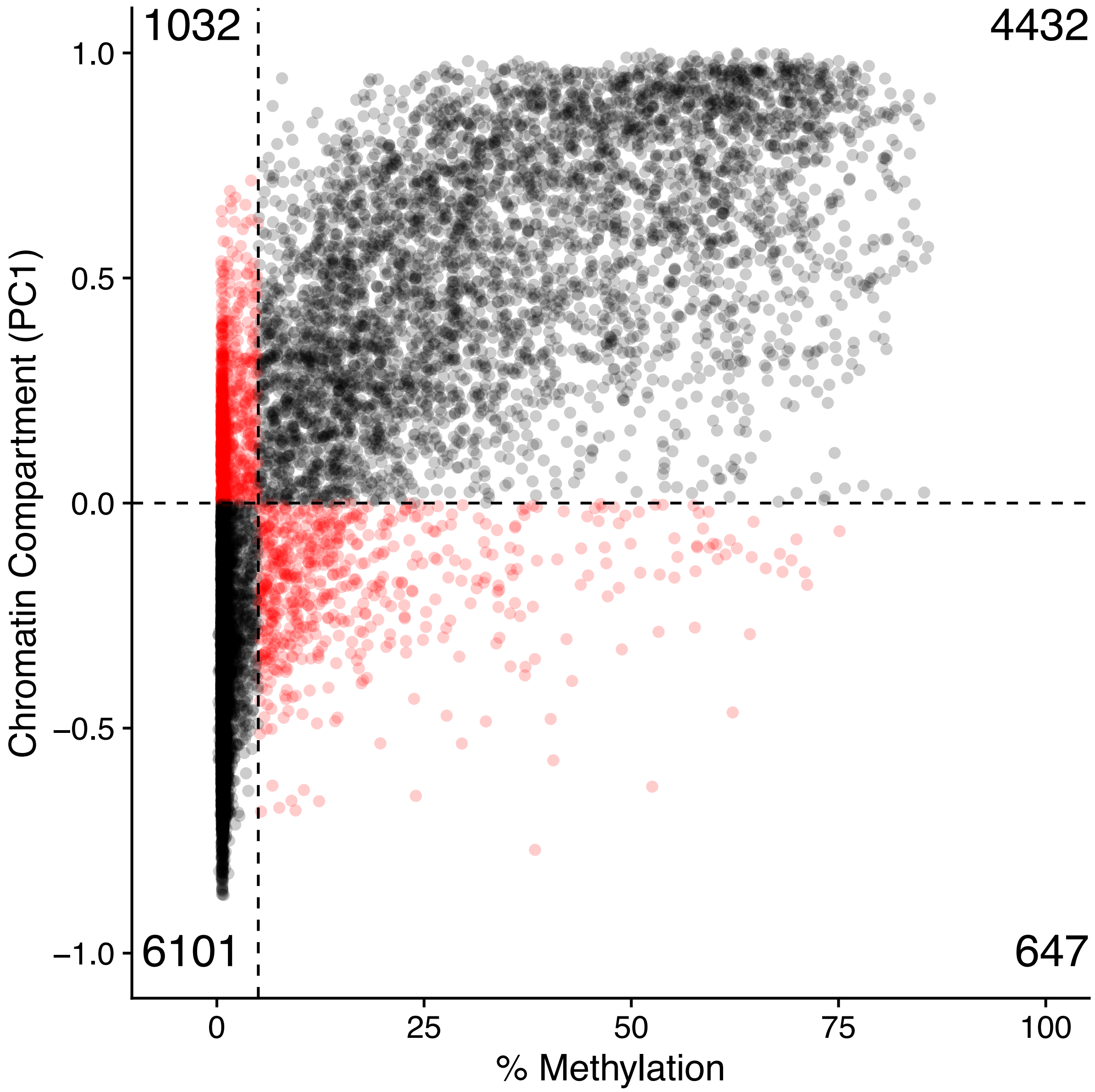


**Figure S18.** Scatter plot of percent DNA methylation versus chromatin compartment score, showing chromatin compartment assignment by each dataset. The vertical dashed line shows the compartment score threshold above which compartments are assigned to euchromatin (PC1 > 0). The horizontal dashed line shows the percent DNA methylation value above which compartments are assigned to euchromatin (% methylation > 5%). Each point is a 100kb bin, with points in red being bins assigned to different chromatin compartments between methods. Numbers in each corner show the number of bins in each quadrant.

### Supplementary Methods

*RNA-seq quality control*

**Table S2.** Summary of RNA-seq samples, sequencing and mapping. See table S6 for coordinates of sampling localities.

| ID | Sample Date | Locality | Host | Morph | Sex | Raw Read Pairs (x$10^6$) | Trimmed Read Pairs (x$10^6$) | Mapped Read Pairs (x$10^6$) |
|---|---|---|---|---|---|---|---|---|
| 17_0003 | 2017.4.25 | N1 | A | G | F | 7.41 | 6.71 | 2.64 |
| 17_0006 | 2017.4.25 | N1 | C | G | F | 8.31 | 7.6 | 3.2 |
| 17_0012 | 2017.4.25 | FH | A | S | F | 7.42 | 6.7 | 2.78 |
| 17_0015 | 2017.4.25 | FH | A | S | F | 8.16 | 7.44 | 2.98 |
| 17_0019 | 2017.4.25 | L | A | S | F | 7.07 | 6.46 | 2.97 |
| 17_0043 | 2017.4.25 | HV | A | G | F | 7.45 | 6.67 | 5.5 |
| 17_0045 | 2017.4.25 | HV | A | S | F | 7.02 | 6.33 | 2.38 |
| 17_0049 | 2017.4.25 | HV | C | S | F | 7.37 | 6.43 | 5.34 |
| 17_0051 | 2017.4.25 | HV | C | M | F | 6.69 | 6.06 | 2.45 |
| 17_0057 | 2017.4.25 | SCN | A | S | F | 7.96 | 7.14 | 3.35 |
| 17_0062 | 2017.4.25 | SC | C | G | F | 8.54 | 7.78 | 6.08 |
| 17_0065 | 2017.4.25 | SC | C | G | F | 6.77 | 6.11 | 2.52 |
| 17_0067 | 2017.4.25 | OUT | A | G | F | 6.1 | 5.47 | 4.65 |
| 17_0070 | 2017.4.25 | OUT | A | G | F | 6.76 | 6.05 | 5.12 |
| 17_0074 | 2017.4.25 | OUT | C | G | F | 7.95 | 7.17 | 0.98 |
| 17_0075 | 2017.4.25 | OUT | C | S | F | 5.91 | 5.4 | 4.46 |
| 17_0081 | 2017.4.25 | PR | C | G | F | 5.28 | 4.74 | 0.84 |
| 17_0082 | 2017.4.25 | BT | A | G | F | 5.2 | 4.71 | 0.62 |
| mean | | | | | | 7.08 | 6.39 | 3.27 |
| s.d. | | | | | | 0.98 | 0.9 | 1.63 |

*Methyl-seq quality control*

**Table S3.** Summary of Methyl-seq samples, sequencing and mapping. CpG coverage calculated as the total number of CpG site bases covered by a read. % methylation calculated at the total genome-wide cytosine count at CpG sites divided by CpG coverage. See Table S6 for coordinates of sampling localities. Local. = locality, Methyl. = methylation.

| ID | Sample Date | Local. | Host | Morph | Sex | Raw Read Pairs ($x10^6$) | Trimmed Read Pairs ($x10^6$) | Mapped Read Pairs ($x10^6$) | CpG coverage ($x10^6$) | % Methyl. |
|---|---|---|---|---|---|---|---|---|---|---|
| 17_0003 | 2017.4.2 | N1 | A | G | F | 40.3 | 39.79 | 18.46 | 95.04 | 14.2 |
| 17_0005 | 2017.4.2 | N1 | A | G | F | 34.02 | 33.3 | 15.3 | 82.69 | 13.5 |
| 17_0006 | 2017.4.2 | N1 | C | G | F | 46.24 | 45.6 | 21.38 | 108.7 | 14 |
| 17_0009 | 2017.4.2 | N1 | C | M | F | 48.96 | 48.28 | 21.42 | 107.51 | 14.5 |
| 17_0012 | 2017.4.2 | FH | A | S | F | 45.98 | 45.35 | 21.37 | 109.67 | 14.3 |
| 17_0015 | 2017.4.2 | FH | A | S | F | 35.5 | 34.69 | 16.38 | 89.04 | 13.7 |
| 17_0018 | 2017.4.2 | L | A | S | F | 33.99 | 33.47 | 13.95 | 75.76 | 14.9 |
| 17_0019 | 2017.4.2 | L | A | S | F | 45.26 | 44.57 | 21.15 | 107.41 | 14.5 |
| 17_0043 | 2017.4.2 | HV | A | G | F | 33.33 | 32.92 | 15.73 | 78.1 | 13.2 |
| 17_0045 | 2017.4.2 | HV | A | S | F | 48.58 | 47.74 | 22.33 | 109.91 | 15 |
| 17_0049 | 2017.4.2 | HV | C | S | F | 38.02 | 37.52 | 17.57 | 89.32 | 12.5 |
| 17_0051 | 2017.4.2 | HV | C | M | F | 31.6 | 31.16 | 15.12 | 76.7 | 13.1 |
| 17_0057 | 2017.4.2 | SCN | A | S | F | 40.41 | 39.88 | 19.5 | 98.96 | 13 |
| 17_0058 | 2017.4.2 | SCN | A | S | F | 37.6 | 36.78 | 15.98 | 88.43 | 14.7 |
| 17_0062 | 2017.4.2 | SC | C | G | F | 37.25 | 36.72 | 17.6 | 93.88 | 13.4 |
| 17_0065 | 2017.4.2 | SC | C | G | F | 33.19 | 32.67 | 15.37 | 78.71 | 12.5 |
| 17_0067 | 2017.4.2 | OUT | A | G | F | 40.81 | 40.3 | 19.04 | 100.78 | 14.2 |
| 17_0070 | 2017.4.2 | OUT | A | G | F | 31.85 | 31.43 | 15.45 | 79.17 | 13.5 |
| 17_0074 | 2017.4.2 | OUT | C | G | F | 46.99 | 46.34 | 22.07 | 112.06 | 14.4 |
| 17_0075 | 2017.4.2 | OUT | C | S | F | 42.53 | 41.86 | 20.21 | 104.08 | 14.1 |
| 17_0077 | 2017.4.2 | PR | C | G | F | 42.82 | 42.23 | 18.75 | 98.09 | 13.1 |
| 17_0081 | 2017.4.2 | PR | C | G | F | 43.8 | 42.86 | 20.54 | 112.05 | 13.6 |
| 17_0082 | 2017.4.2 | BT | A | G | F | 46.28 | 45.6 | 20.68 | 109.45 | 13.6 |
| 17_0086 | 2017.4.2 | BT | A | G | F | 48.93 | 48.27 | 21.93 | 109.9 | 13.3 |
| 23_0597 | 2023.5.8 | PR | C | G | F | 121.99 | 117.77 | 66.7 | 383.08 | 11.2 |
| 23_0599 | 2023.5.8 | PR | C | G | F | 93.58 | 90.7 | 52.07 | 293.68 | 11.1 |
| 23_0601 | 2023.5.8 | PR | C | G | F | 96.02 | 92.65 | 51.92 | 299.56 | 11.1 |
| 23_0602 | 2023.5.8 | PR | C | G | F | 94.71 | 91.87 | 56.23 | 329.14 | 11.1 |
| 23_0608 | 2023.5.8 | PR | C | G | F | 131.15 | 126.82 | 66.7 | 386.27 | 11.1 |
| 23_0613 | 2023.5.8 | PR | C | G | F | 98.35 | 95.23 | 54.43 | 303.78 | 10.5 |
| 23_0622 | 2023.5.8 | PR | C | M | F | 111.16 | 107.4 | 51.84 | 295.81 | 11.7 |
| 23_0631 | 2023.5.8 | PR | C | G | F | 110.52 | 106.95 | 59.72 | 331.29 | 11.2 |
| 23_0635 | 2023.5.8 | PR | C | G | F | 101.41 | 98.19 | 59.05 | 324.73 | 11.1 |
| 23_0636 | 2023.5.8 | PR | C | G | F | 115.32 | 111.33 | 58.89 | 337.21 | 11.7 |
| mean 2017 | | | | | | 40.59 | 39.97 | 18.64 | 96.48 | 13.8 |
| mean 2023 | | | | | | 107.42 | 103.89 | 57.76 | 328.45 | 11.18 |
| mean total | | | | | | 60.25 | 58.77 | 30.14 | 164.7 | 13.03 |
| s.d. 2017 | | | | | | 5.83 | 5.76 | 2.68 | 12.87 | 0.71 |
| s.d. 2023 | | | | | | 12.75 | 12.24 | 5.59 | 33.57 | 0.34 |
| s.d. total | | | | | | 60.25 | 58.77 | 30.14 | 164.7 | 1.36 |

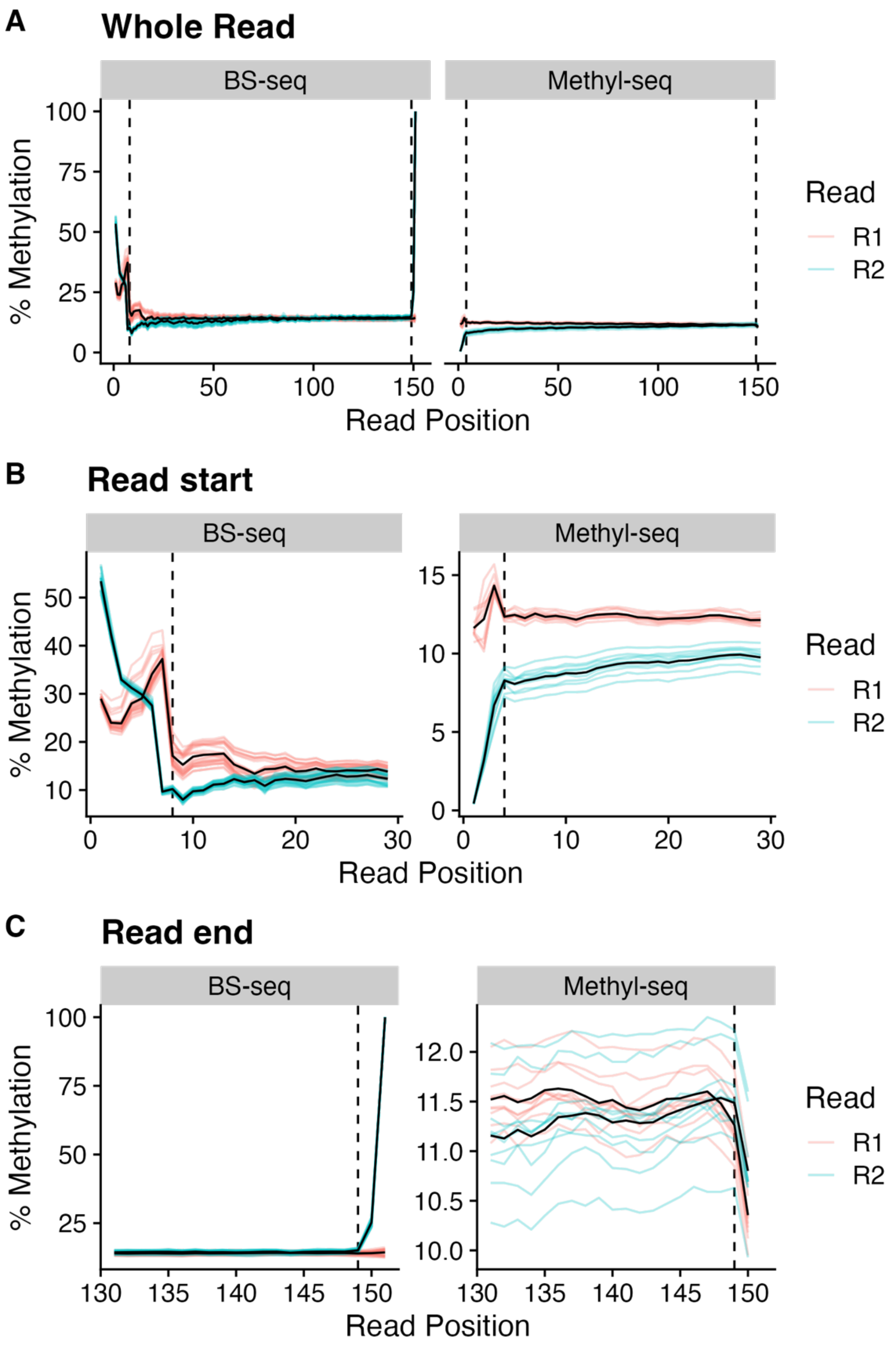


**Figure S19.** Bias in % methylation by read position. Lighter coloured lines are the average for each read for each individual and the black lines are the average among individuals for each read. (A) Overall bias. (B) Bias at the start of reads. Positions to the left of vertical dashed line were filtered out. (C) Bias at the end of reads. Position to the right of the vertical dashed line were filtered out.

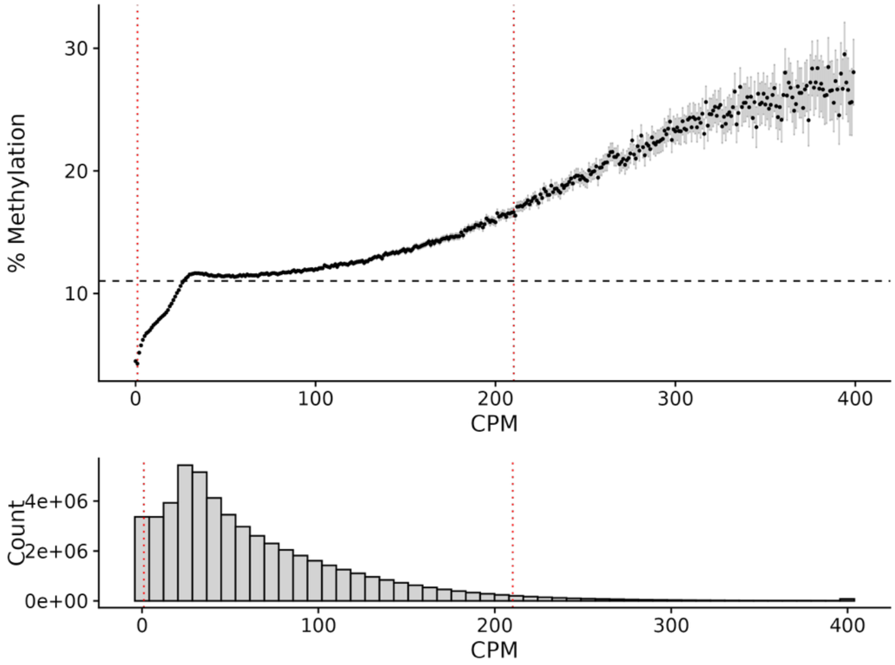


**Figure S20.** Bias in % methylation of CpG sites by count per million reads (CPM). Points are the average % methylation for all CpG sites with a given CPM (rounded to the nearest whole number). All sites with CPM > 400 were reduced to CPM=400 for visualization but not for filtering. Error bars are 95% CI of the mean. The horizontal dashed line is the total average % methylation and the red vertical dotted lines indicate the 95% quantiles for CPM used for filtering.

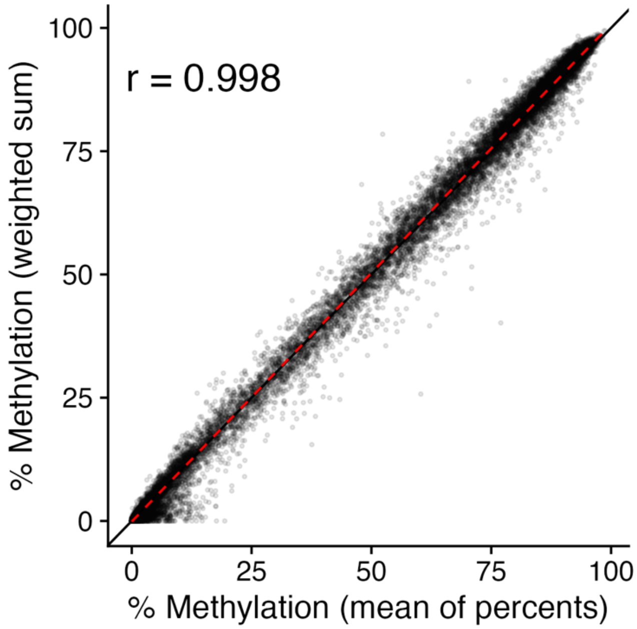


**Figure S21** Correlation between % methylation calculated as the mean of the percent for each individual and % methylation calculated as the sum of C and T across individuals. Each point is an individual CpG site. Only sites with at least five reads for all individuals were included. The black line indicates 1:1. The red dotted line gives the linear line of best fit.

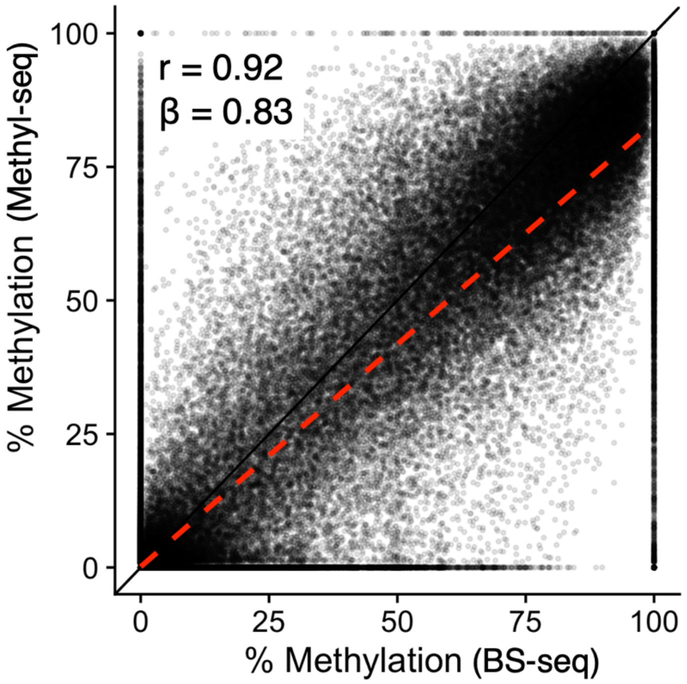


**Figure S22.** Correlation in % methylation at CpG sites between years. % methylation was calculated by summing count per million reads (CPM) of C and T over all individuals within a year. Each point is a CpG site. The black line indicates 1:1. The red dotted line gives the linear line of best fit.

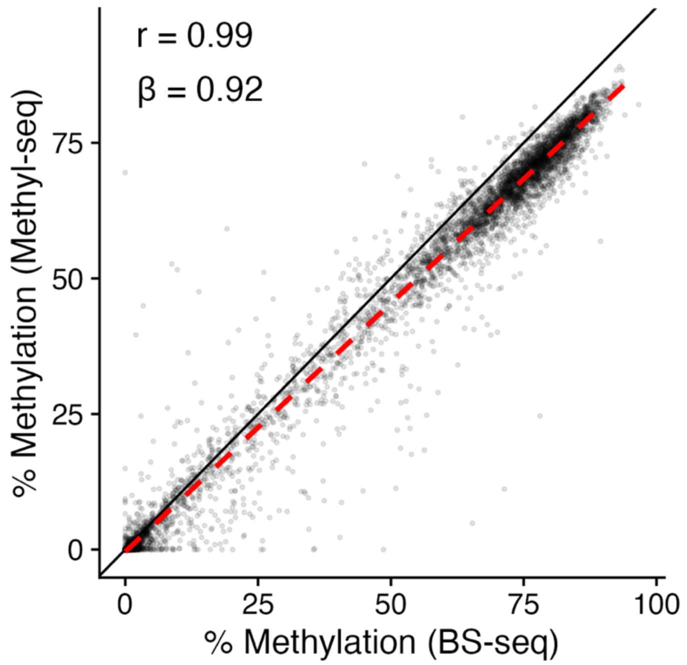


**Figure S23.** Correlation in % methylation in gene bodies between years. % methylation was calculated by summing count per million reads (CPM) counts among CpG sites within each gene, then summing over all individuals within a year. Each point is a gene. The black line indicates 1:1. The red dotted line gives the linear line of best fit.

*ATAC-seq quality control*

**Table S4.** Summary of ATAC-seq samples, sequencing and mapping. FRiP = fraction of reads in peaks, TSS = transcription start site. TSS enrichment calculated by taking the ratio of the highest fragment count within the 500 bp upstream of the TSS to the fragment count of 500 bp regions 1500 bp from the TSS. See Table S6 for coordinates of sampling localities.

| ID | Sample Date | Locality | Host | Morph | Sex | Raw Read Pairs (x10$^6$) | Trimmed Read Pairs (x10$^6$) | Mapped Read Pairs (x10$^6$) | Peaks (x10$^3$) | FRiP score | TSS Enrichment |
|---|---|---|---|---|---|---|---|---|---|---|---|
| 15_0694 | 2015.4.18 | FH | A | G | F | 17.88 | 17.61 | 10.12 | 41.1 | 0.17 | 6.69 |
| 15_0724 | 2015.4.18 | L | A | G | F | 58.1 | 56.77 | 22.44 | 81.84 | 0.18 | 5.88 |
| 15_0768 | 2015.4.18 | HV | A | G | F | 55.72 | 52.63 | 5.17 | 4.05 | 0.11 | 2.13 |
| 15_0769 | 2015.4.18 | HV | A | G | F | 53.85 | 52.41 | 22.54 | 62.65 | 0.17 | 4.16 |
| 15_0775 | 2015.4.18 | HV | C | G | F | 53.19 | 41.64 | 4.69 | 3.96 | 0.12 | 2.38 |
| 15_0778 | 2015.4.18 | HV | C | G | F | 54.54 | 53.45 | 19.97 | 51.92 | 0.18 | 3.99 |
| 21_0010 | 2021.5.30 | PR | C | G | F | 30 | 29.68 | 14.27 | 44.49 | 0.14 | 4.65 |
| 21_0011 | 2021.5.30 | PR | C | G | F | 30 | 29.59 | 14.69 | 55.77 | 0.19 | 6.98 |
| 21_0018 | 2021.5.30 | L | A | S | F | 30 | 29.45 | 11.58 | 88.98 | 0.21 | 9.78 |
| 21_0019 | 2021.5.30 | L | A | S | F | 30 | 29.47 | 14.67 | 104.26 | 0.2 | 7.66 |
| mean | | | | | | 41.33 | 39.27 | 28.86 | 53.9 | 0.17 | 5.43 |
| s.d. | | | | | | 15 | 13.79 | 13.09 | 33.1 | 0.03 | 2.42 |

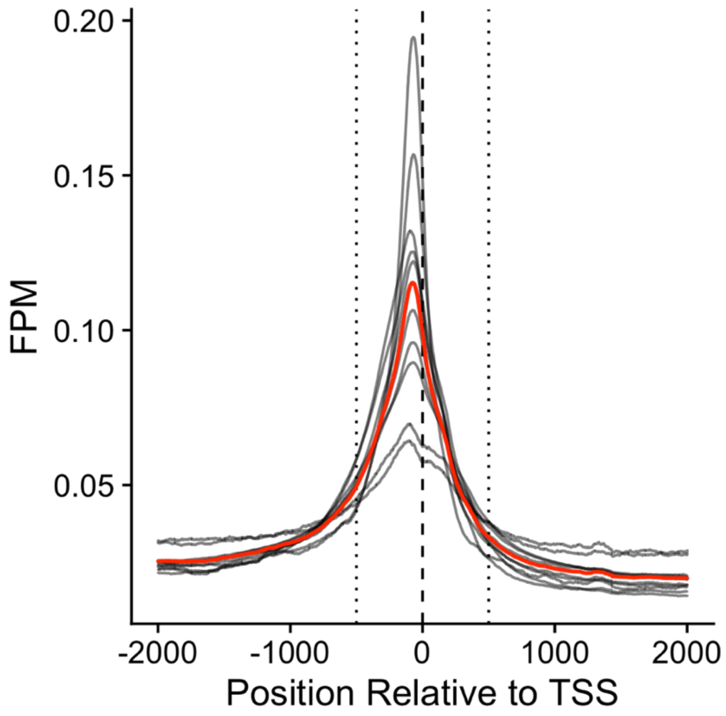


**Figure S24.** Enrichment of ATAC-seq fragments per million fragments (FPM) at transcription start sites (TSSs). Black lines are values for an individual, averaged over all TSSs. The red line is the average among individuals. The vertical dashed line indicates the TSS. The vertical dotted lines indicate ± 500 bp from the TSS. The 500 bp to the left (upstream) of the TSS is the putative promoter region whereas the 500 bp to the right (downstream) of the TSS is the putative 'head' region of the gene.

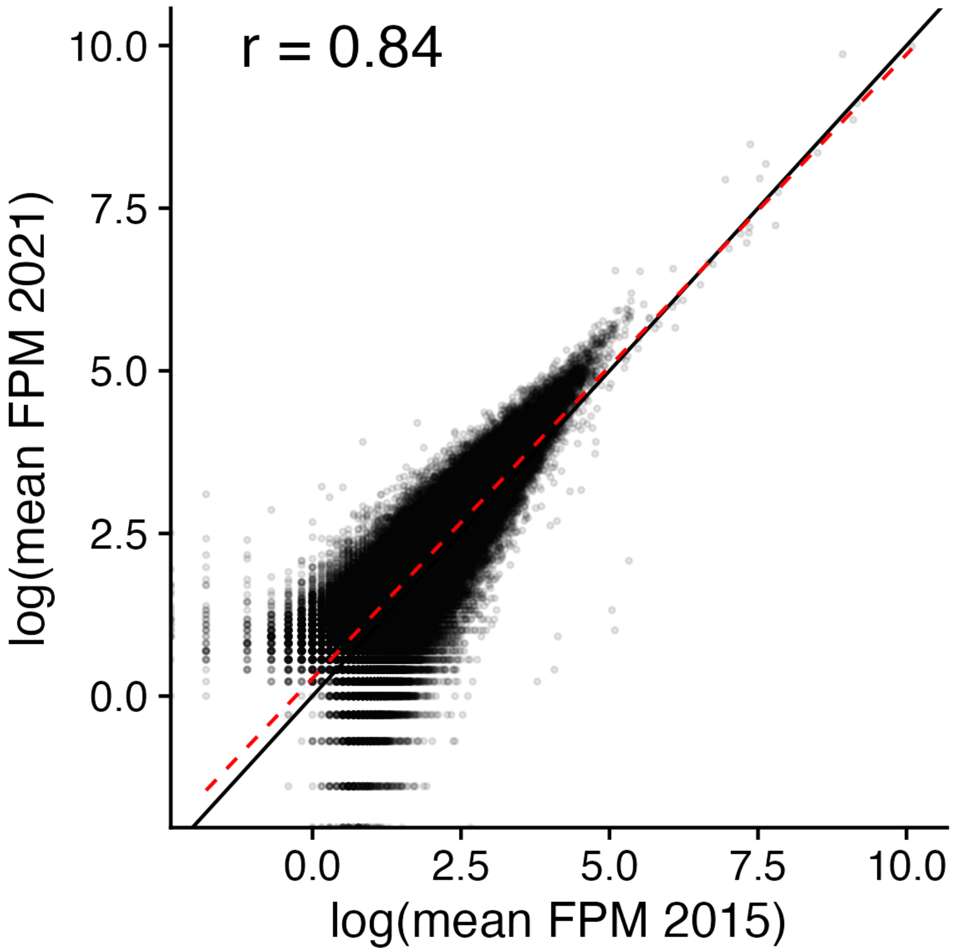


**Figure S25.** Correlation between batches for fragments per million fragments (FPM) in open chromatin peaks. Each point is an individual open chromatin peak. The black line indicates 1:1. The red dotted line gives the linear line of best fit.

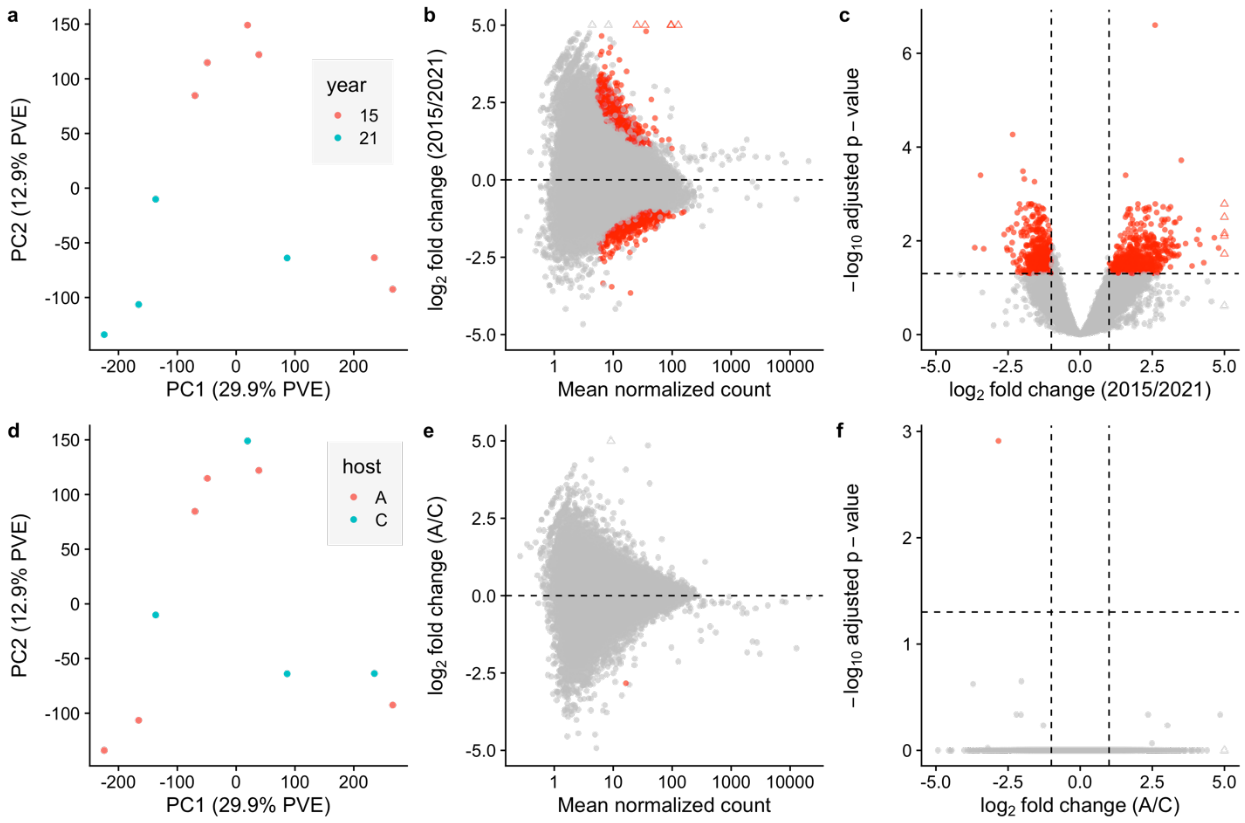


**Figure S26.** Difference in openness of chromatin peaks between sample batches (a-c) and host-plant ecotypes (d-f). All plots are of fragments counts within consensus open chromatin peaks, from ATAC-seq reads down sampled for even coverage among individuals. (a,d) PCs calculated from log-transformed fragment counts. Each point is an individual *T. cristinae*. A = *Adenostoma fasculatum*, C = *Ceanothus spinosus*. (b,e) MA plots. (c,f) Volcano plots. (b,c,e,f) Points are individual open chromatin peaks, with red points indicating peaks which significantly differ between groups after correction for multiple testing. Triangular points are those with $|\log_2$ fold change$| > 5$.

*Omni-C / Hi-C quality control*

**Table S5.** Summary of Hi-C samples, sequencing and mapping. See Table S6 for coordinates of sampling localities.

| ID | Sample Date | Locality | Host | Morph | Sex | Raw Reads ($x10^6$) | Trimmed Reads ($x10^6$) | Mapped Reads ($x10^6$) | # Short-range Contacts ($x10^6$) | # Long-range Contacts ($x10^6$) |
|---|---|---|---|---|---|---|---|---|---|---|
| CEN4119 | 2023.05.15 | VP | A | S | F | 182.21 | 180.95 | 75.38 | 18.24 | 50.99 |
| CEN4280 | 2023.05.15 | VP | A | G | F | 146.56 | 145.37 | 56.82 | 17.41 | 34.05 |
| CEN1559 | 2019 | PRN | C | G | F | 260.57 | 259.69 | 129.39 | 93.04 | 32.61 |
| mean | | | | | | 196.44 | 195.34 | 87.2 | 42.9 | 39.22 |
| s.d. | | | | | | 58.32 | 58.5 | 37.7 | 43.43 | 10.22 |

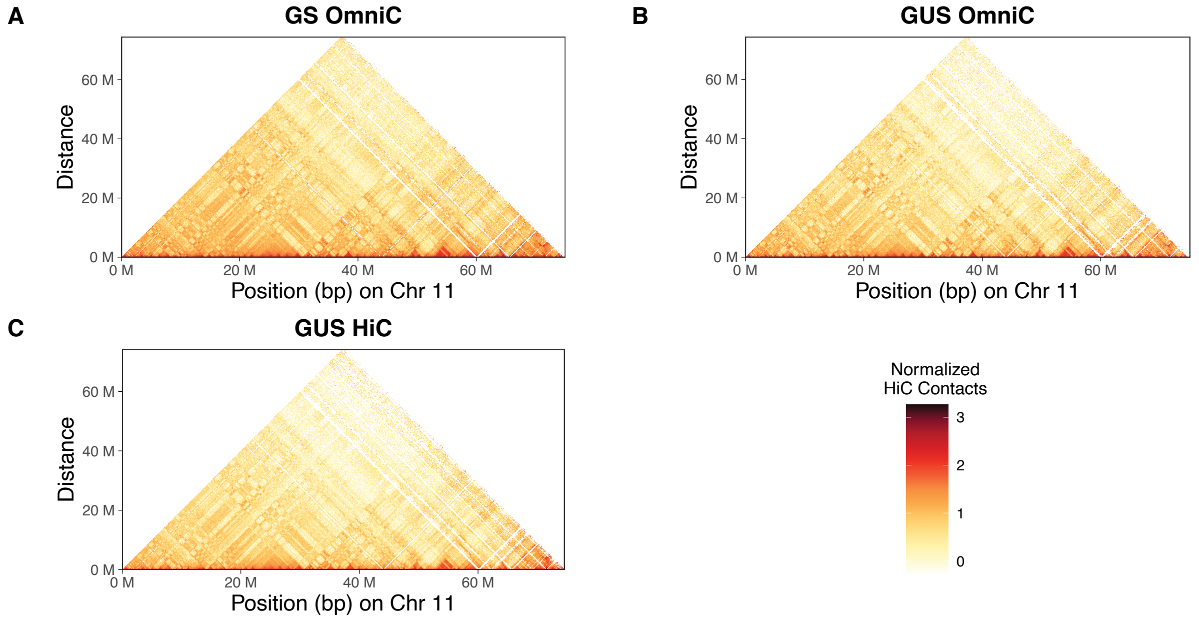


**Figure S27.** Hi-C contact maps on chromosome 11, as an example of concordance in patterns of Hi-C contacts between samples.

*Sampling sites*

**Table S6.** Latitude and longitude of sampling sites.

| Locality | Latitude | Longitude |
|---|---|---|
| BT | 34.536 | -119.862 |
| FH | 34.518 | -119.801 |
| HV | 34.488 | -119.787 |
| L | 34.509 | -119.796 |
| N1 | 34.517 | -119.797 |
| OUT | 34.532 | -119.843 |
| PR | 34.533 | -119.832 |
| SC | 34.523 | -119.832 |
| SCN | 34.521 | -119.830 |

*Comparative analysis of* Apis mellifera

**Table S7.** Summary of *Apis mellifera* RNA-seq samples, sequencing and mapping.

| Study | Accession | Name | Raw Read Pairs ($x10^6$) | Trimmed Read Pairs ($x10^6$) | Mapped Read Pairs ($x10^6$) |
|---|---|---|---|---|---|
| Jin et al. 2023 | SRR18219008 | Worker 1 | 24.44 | 24.43 | 22.81 |
| Jin et al. 2023 | SRR18219007 | Worker 2 | 22.82 | 22.81 | 21.53 |
| Jin et al. 2023 | SRR18219006 | Worker 3 | 21.59 | 21.58 | 19.9 |
| Lowe et al. 2022 | SRR19861408 | Worker 1 | 17.55 | 17.35 | 15.68 |
| Lowe et al. 2022 | SRR19861409 | Worker 2 | 28.82 | 28.75 | 26.07 |
| mean | | | 23.04 | 22.98 | 21.2 |
| s.d. | | | 4.11 | 4.16 | 3.83 |

**Table S8.** Summary of *Apis mellifera* BS-seq samples, sequencing and mapping. CpG coverage calculated as the total number of CpG site bases covered by a read. % methylation calculated at the total genome-wide cytosine count at CpG sites divided by CpG coverage.

| Study | Accession | Name | Raw Read Pairs ($x10^6$) | Trimmed Read Pairs ($x10^6$) | Mapped Read Pairs ($x10^6$) | CpG coverage ($x10^6$) | % Methylation |
|---|---|---|---|---|---|---|---|
| Rasmussen et al. 2021 | SRR14037562 | Workers A1 | 18.09 | 18.02 | 10.84 | 123.48 | 0.9 |
| Rasmussen et al. 2021 | SRR14037563 | Workers A2 | 27.82 | 27.71 | 17.25 | 138.02 | 1 |
| Rasmussen et al. 2021 | SRR14037567 | Workers B1 | 23.88 | 23.78 | 12.56 | 118.2 | 1 |
| Rasmussen et al. 2021 | SRR14037568 | Workers B2 | 22.96 | 22.9 | 11.65 | 95.69 | 1.1 |
| Rasmussen et al. 2021 | SRR14037569 | Workers B3 | 27.4 | 27.35 | 14.45 | 111.08 | 1.2 |
| mean | | | 24.03 | 23.95 | 13.35 | 117.29 | 1.04 |
| s.d. | | | 3.94 | 3.94 | 2.56 | 15.6 | 0.11 |

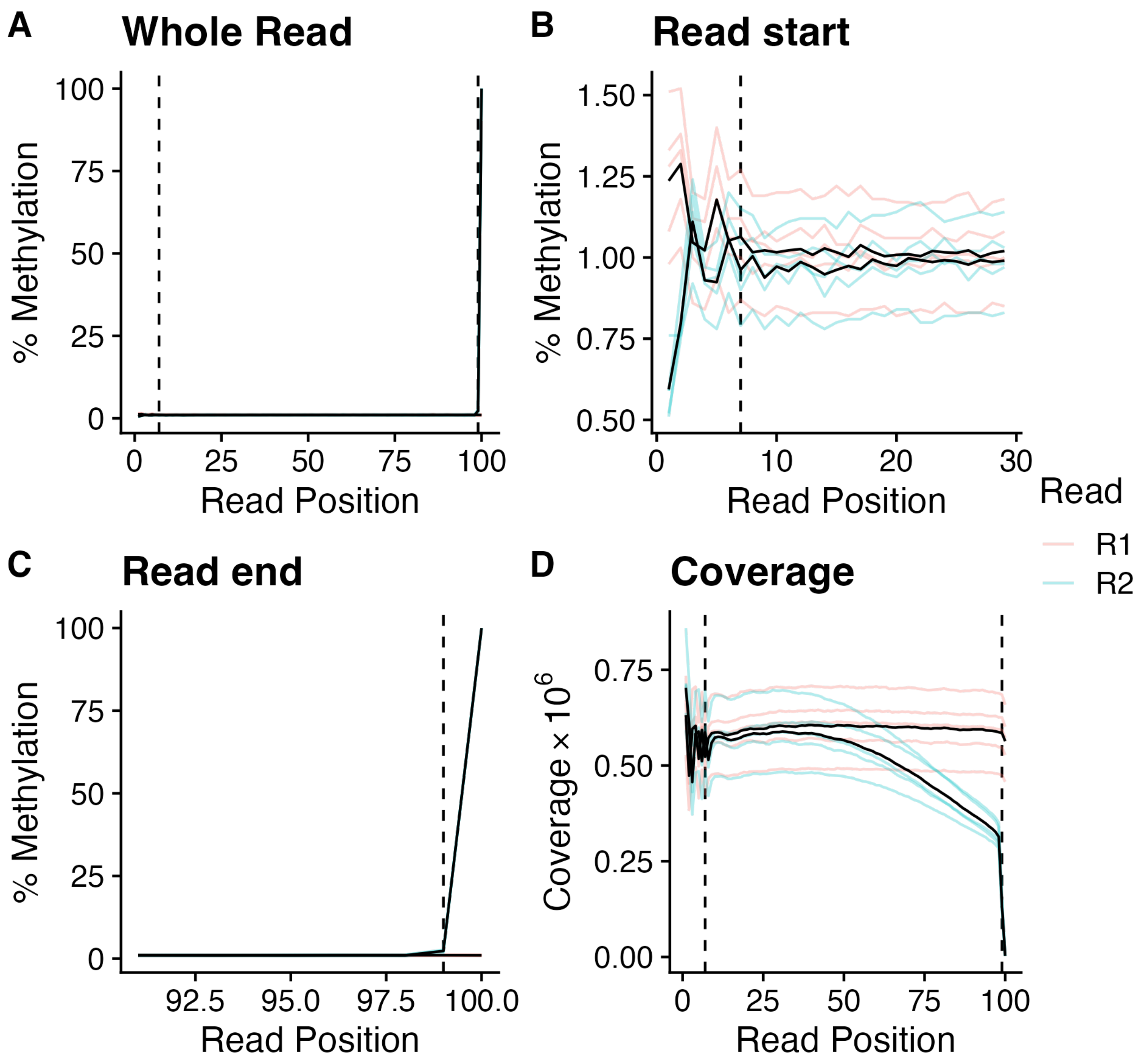


**Figure S28.** Bias in % methylation by read position for *Apis mellifera* samples. Lighter coloured lines are the average for each read for each individual and the black lines are the average among individuals for each read. (A) Overall bias. (B) Bias at the start of reads. Positions to the left of vertical dashed line were filtered out. (C) Bias at the end of reads. Position to the right of the vertical dashed line were filtered out. (D) Coverage per read position

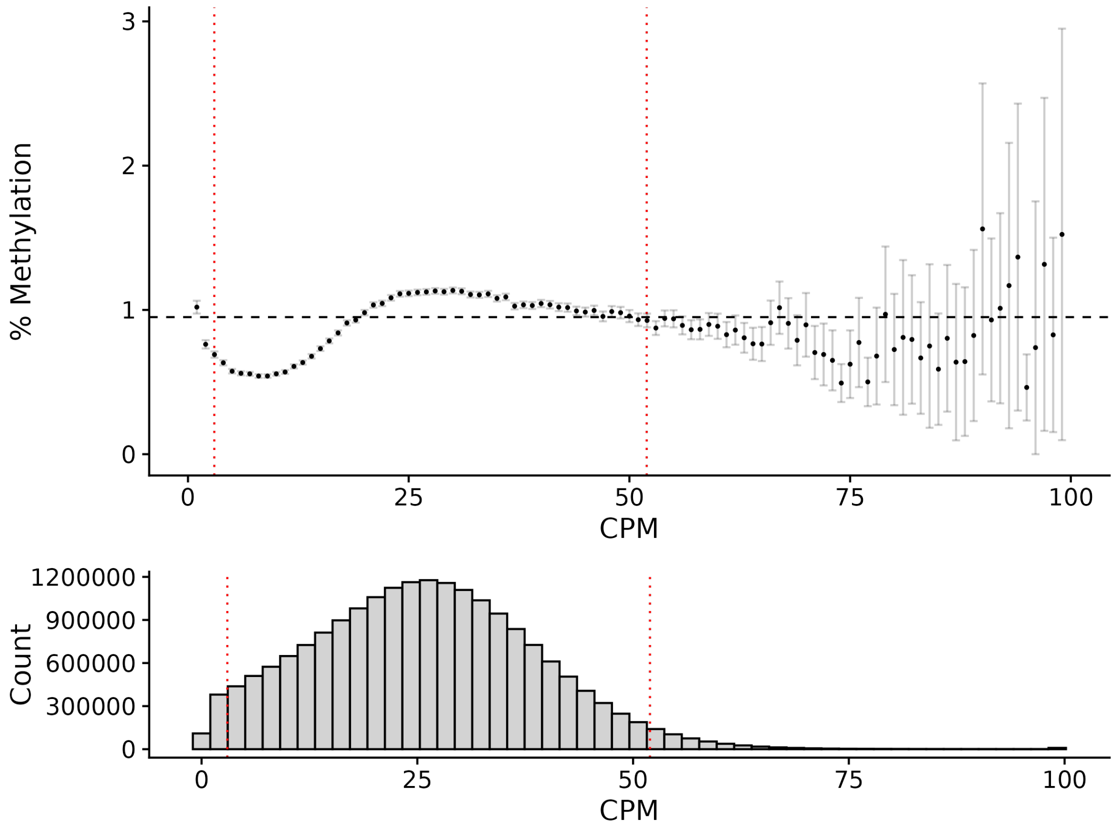


**Figure S29.** Bias in % methylation of CpG sites by count per million reads (CPM) for *Apis mellifera* samples. Points are the average % methylation for all CpG sites with a given CPM (rounded to the nearest whole number). All sites with CPM > 400 were reduced to CPM=400 for visualization but not for filtering. Error bars are 95% CI of the mean. The horizontal dashed line is the total average % methylation and the red vertical dotted lines indicate the 95% quantiles for CPM used for filtering.

**Table S9.** Summary of *Apis mellifera* ATAC-seq samples, sequencing and mapping. FRiP = fraction of reads in peaks, TSS = transcription start site. TSS enrichment calculated by taking the ratio of the highest fragment count within the 500 bp upstream of the TSS to the fragment count of 500 bp regions 1500 bp from the TSS.

| **Study** | **Accession** | **Name** | **Raw Read Pairs ($x10^6$)** | **Trimmed Read Pairs ($x10^6$)** | **Mapped Read Pairs ($x10^6$)** | **Peaks ($x10^3$)** | **FRiP score** | **TSS Enrichment** |
|---|---|---|---|---|---|---|---|---|
| Lowe et al. 2022 | SRR19866069 | Worker 1 | 24.16 | 23.97 | 9.23 | 31.39 | 0.5 | 2.95 |
| Lowe et al. 2022 | SRR19866070 | Worker 2 | 26.1 | 25.9 | 9.11 | 29.04 | 0.48 | 2.71 |
| mean | | | 25.13 | 24.93 | 9.17 | 30.21 | 0.49 | 2.83 |
| s.d. | | | 1.38 | 1.37 | 0.09 | 1.66 | 0.01 | 0.17 |

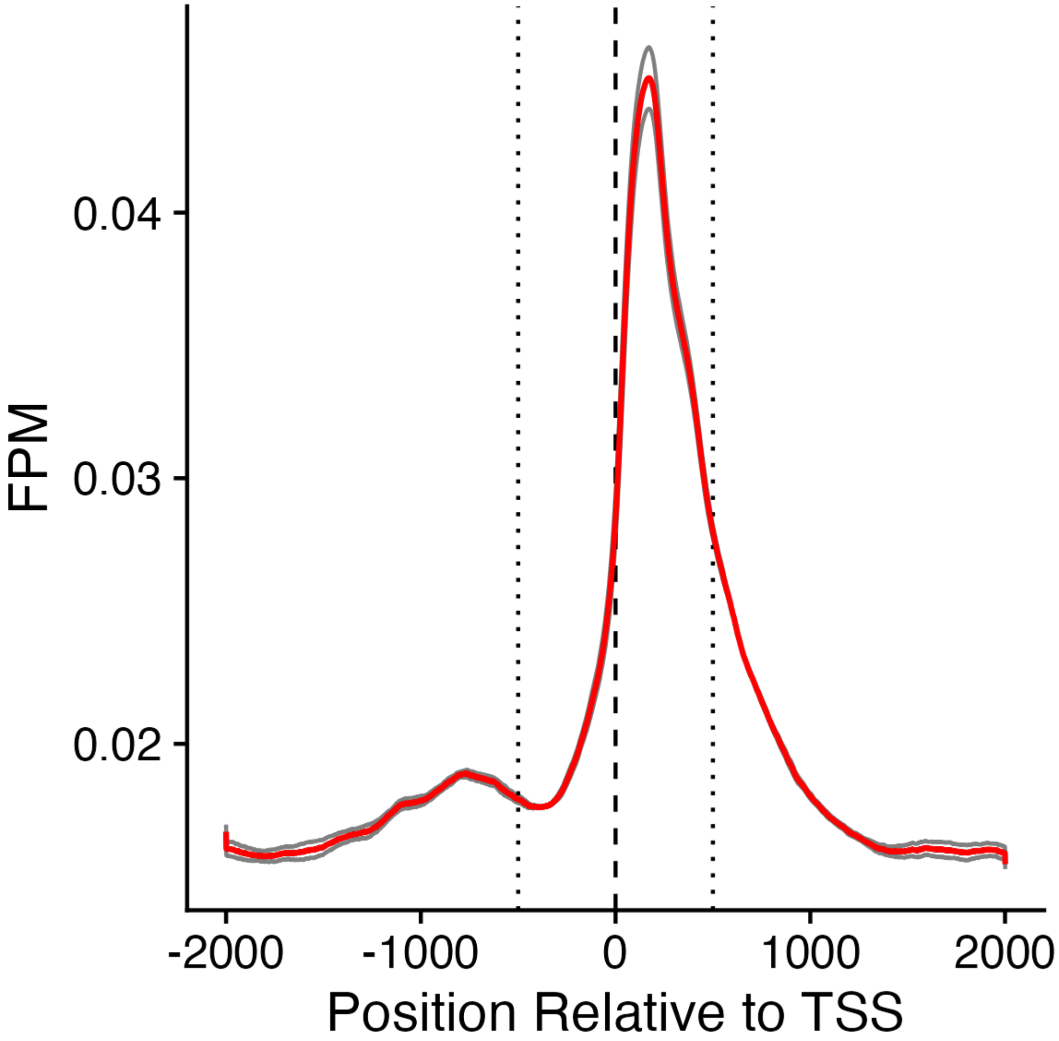


**Figure S30.** Enrichment of ATAC-seq fragments per million fragments (FPM) at transcription start sites (TSSs) for *Apis mellifera* samples. Black lines are values for an individual, averaged over all TSSs. The red line is the average among individuals. The vertical dashed line indicates the TSS. The vertical dotted lines indicate ± 500 bp from the TSS. The 500 bp to the left (upstream) of the TSS is the putative promoter region whereas the 500 bp to the right (downstream) of the TSS is the putative 'head' region of the gene.

**Table S10.** Summary of *Apis mellifera* Hi-C sample, sequencing and mapping.

| Study | Accession | Name | Raw Reads ($x10^6$) | Trimmed Reads ($x10^6$) | Mapped reads ($x10^6$) | # Short-range Contacts ($x10^6$) | # Long-range Contacts ($x10^6$) |
|---|---|---|---|---|---|---|---|
| Jin et al. 2023 | SRR18355687 | Worker | 605.93 | 605.65 | 392.16 | 110.44 | 171.67 |
